# Two polybasic domains of septin9 control septin filaments assembly and Golgi integrity

**DOI:** 10.1101/535039

**Authors:** Mohyeddine Omrane, Amanda S. Camara, Cyntia Taveneau, Nassima Benzoubir, Thibault Tubiana, Jinchao Yu, Raphaël Guérois, Didier Samuel, Bruno Goud, Christian Poüs, Stéphane Bressanelli, Richard C. Garratt, Abdou Rachid Thiam, Ama Gassama-Diagne

**Author notes:** Joint senior and corresponding authors Correspondence to or.

## Abstract

Septins are GTP-binding proteins involved in several membrane remodeling mechanisms. They associate with membranes, presumably by using a polybasic domain (PB1) that interacts with phosphoinositides (PIs). Membrane-bound septins assemble into microscopic structures that regulate membrane shape. How septins exactly interact with PIs, assemble, and shape membranes is weakly understood. Here, we found that septin 9 has a second polybasic domain (PB2) conserved in the human septin family. Similarly to PB1, PB2 binds specifically to PIs, and both domains are critical for septin filament formation. However, septin 9 membrane association does not depend on these PB domains but on putative PB-adjacent amphipathic helices. The presence of the PB domains guarantees the protein enrichment to PI-contained membranes, which is critical for PI-enriched organelles. In particular, we found that septin 9 PB domains control the assembly and functionality of the Golgi apparatus. Our findings bring novel insights into the role of septins in organelle morphology.

**Highlights:**

- Two polybasic domains mediate septin 9 interaction with PIs
- Human septins have amphipathic helices suitable for binding membrane
- Septin 9 polybasic domains mediate septin high order structure formation
- Mutation or depletion of septin polybasic domains induce Golgi fragmentation

## Introduction

Septins form a GTPase protein family found in eukaryotes from yeasts to animals, but absent from higher plants and certain protists (Pan et al., 2007; Nishihama et al.; 2011). In mammals, thirteen septins have been identified and classified into four classes (the septin 2, 3, 6 and 7 subgroups) (Peterson and Petty, 2010). Septins assemble into apolar complexes that are able to form high order structures such as filaments and rings (Weirich et al., 2008). Each septin has at least two interfaces: one interface contains GTP-binding domain motifs, referred to as the G-interface, and the other one contains the N and C termini of the protein, called the NC-interface. Thus, two septin proteins can develop G/G or NC/NC interactions with neighbor septins. Septins thereby form hetero-oligomeric complexes made up of hexameric subunits with the following sequence: (G7NC/NC6G/G2NC/NC2G/G6NC/NC7G) (Sirajuddin et al., 2007). Septin 9 assembles at the extremities of the hexamer to generate an octamer (Kim et al., 2011). This octamer has the NC interface of septin 9 at its ends, i.e. NC9G/G7NC/NC6G/G2NC/NC2G/G6NC/NC7G/G9NC, and is the building block for higher order septin structures (Sirajuddin et al., 2007, Sellin et al., 2011, Kim et al., 2011). During membrane remodeling processes, these structures can act as a diffusion barrier or scaffolds that recruit cytosolic proteins and other cytoskeletal elements such as microtubules or actins (Tanaka-Takiguchi et al., 2009; Fung et al., 2014; Bridges et al., 2016; Mostowy and Cossart, 2012).

Septins bind specifically to phosphoinositides (PIs) via a polybasic domain (PB1) located at the N-terminus of their GTP-binding domain. This interaction with PIs supposedly mediates septin membrane association, which is a determinant factor for the structural and functional features of the protein (Pan et al., 2007; Zhang et al., 1999; Tanaka-Takiguchi et al., 2009; Casamayor and Snyder, 2003). Septins associate with a variety of PIs at different intracellular membranes (Zhang et al., 1999; Akil et al., 2016; Dolat and Spiliotis, 2016; Pagliuso et al., 2016), and control numerous cellular functions such as cytokinesis, cilogenesis, vesicular trafficking, cell polarity and lipid droplet formation (Fung et al., 2014; Oh and Bi, 2011; Song et al., 2016; Balla, 2013; Gassama-Diagne and Payrastre, 2009; Gassama-Diagne et al., 2006; Akil et al., 2016; Schink et al., 2016).

The morphology and positioning of intracellular organelles such as the Golgi and the endoplasmic reticulum (ER) are critical for the proper transport and delivery of vesicles to, maintain cell polarity, tissue homeostasis and functions (Lavieu et al., 2014; de Forges et al., 2012; van Bergeijk et al., 2016). Such morphological arrangements are often ensured by cytoskeleton factors such as microtubules and actins, which are in part recruited by septins (Tanaka-Takiguchi et al., 2009; Fung et al., 2014; Bridges et al., 2016; Mostowy and Cossart, 2012). However, whether septins directly affect organelle morphology and function is poorly understood (Gurel et al., 2014; Weirich et al., 2008).

Here, by using septin 9 crystal structures and molecular dynamics simulations (MD) of the septin 9 monomer and dimer, we identified the existence of a second polybasic domain (PB2). The deletion of PB2 phenocopied PB1 deletion in reducing the binding capacity of septin 9 to PI lipids and impairing the formation of septin filamented structures, hereafter referred simply to as filaments. However, *in vitro* flotation assays revealed that the PB domains are not required for septin 9 membrane binding but proffer specificity to PI-containing membranes. These findings prompted us to identify amphipathic helices that are adjacent to the PB domains and possibly mediate the physical association of septin 9 with membranes. We studied the importance of the PB domains on organelles and found their critical role on Golgi assembly and functionality.

## Results

### Septins have a second polybasic domain PB2 that forms with PB1 a basic cluster at the NC interface

Septins bind to PI lipids via a polybasic domain (PB1) located at the N-terminus of their GTP-binding domain (Zhang et al., 1999; Pan et al., 2007). However, we recently found that the deletion of PB1 in septin 9 reduces but does not abolish the interaction between septin 9 and monophosphorylated PIs (Akil et al., 2016). This observation prompted us to look for the presence of additional PI-interacting domains. We aligned the sequences of septin 9 and other human septins and identified a second motif enriched in basic amino acids (aa399-402 of human septin 9 isoform 1; 586 residues) (Figure 1A). This second polybasic domain, which we termed PB2, contains a variable number of basic amino acids (2 to 4) but is conserved in all human septins (Figure 1A).

**Figure 1:**
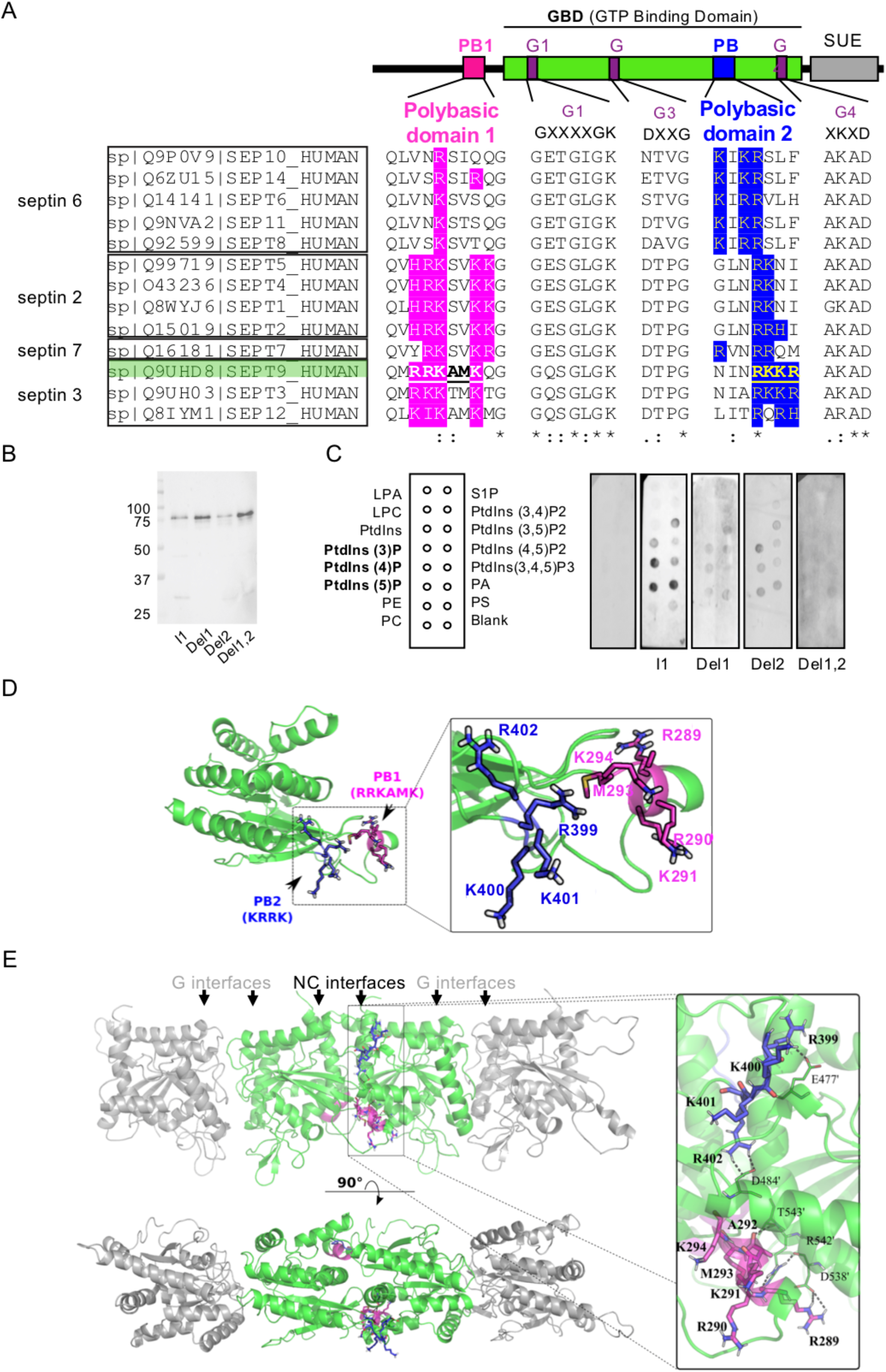
Septin 9 and human septins have two polybasic domains. A. Above: schematic representation of the organization of septin domains. Below: Multiple alignments of human septins: the polybasic domain 1 (PB1), and polybasic domain 2 (PB2), are highlighted in magenta and blue, respectively. Human septin subgroups are shown in boxes. B. Western blot of purified septin 9_i1 and septin 9_del1, septin 9_del2 and septin 9_del1,2. C. PIP strip overlay assay: PIP strips were incubated with either purified septin 9_i1 (I1), septin 9_del1 (Del1), septin 9_del2 (Del2) or septin 9_del1,2 (Del1,2) proteins at 0.5 0.5 µg.ml^-1^ or with the V5 tag peptide as a negative control, and analyzed using the anti-V5 antibody. LPA (lysophosphatidic acid), LPC (lysophosphocholine), PtdIns (phosphatidylinositol), PtdIns(3)P, PtdIns(4)P, PtdIns(5)P, PtdIns(3,4)P2, PtdIns(3,5)P2, PtdIns(4,5)P2, PtdIns(3,4,5)P3, PA (phosphatidic acid), PS (phosphatidylserine), PE (phosphatidylethanolamine), PC (phosphatidylcholine) and S1P (sphingosine 1-phosphate). D. Model of septin 9 based on the crystal structure (PDB code 5cyp) showing PB1 and PB2. **E.** Model of the septin G9NC/NC9G complex using the simulated dimer of septin 9 at the NC interface and based on the symmetry operations of the crystallographic structure (PDB code 5cyp). The two molecules of septin 9 on either side of the NC interface are in green, and their encompassed PB1 and PB2 are presented in magenta and blue, respectively. The rectangle indicates PB1 and PB2 shown at a higher magnification on the right. The residues for PB1 and PB2 are labeled and outlined in black. Dashed black lines indicate the interaction between PBs and neighboring septin 9 residues.

We next generated and purified a PB2-deleted mutant (septin 9_del2), a PB1-deleted mutant (septin 9_del1)(Akil et al., 2016), and a mutant lacking both PB1 and PB2 (septin 9_del1,2) (Figure S1A). These proteins displayed band profiles similar to septin 9_i1 (Figure 1B, Figure S1B), which was in a monomeric form based on migration on a native gel (Figure S1C). We then used a PIP strip overlay assay to determine the affinity of septin 9_i1 and its mutant forms with different phospholipid headgroups. As expected, we found a specific interaction between septin 9_i1 and PtdIns monophosphates (Figure 1C). The interaction signal with PIs was decreased in septin 9_del1 and septin_del2, and it was almost abolished in septin 9_del1,2 (Figure 1C). This result supports that both PB domains can mediate the interaction of septin 9 with phosphoinositides.

To study the involvement of PB2 in the structural organization of septin 9, we opted for a MD simulation approach. We used the most resolved septin 9 crystal structure (aa 293-564), PDB code 5cyp. In this structure, the missing residues and side chains were added and completed by amino acids from 276 to 292 (see methods procedure), which included the ones of PB1. Regardless of the initial folded state of these added residues, we found one single final equilibrium conformation of the protein where the N-terminal region was pre-folded into an α-helix around PB1 (Figure 1D, Figure S1D). This equilibrated monomer was then superposed to the septin 9 crystal structure (PDB code 5cyp) to build a tetramer that contains the NC-NC interface (Figure 1E). At this interface, PB2 and PB1 appeared to make numerous salt bridges between septin monomers (Figure 1E, black dots in the inset), we found optimal distances between amino acids of the PB domains and those of the adjacent septin. These interactions involved the residues R399 and R402 of PB2, respectively interacting with E477’ and D484’, and the residue R289 of PB1 interacting with D538’. The main chain atoms of R289 and A292 in PB1 made hydrogen bonds with R542’ and T543’ of the neighbor septin (Figure 1E). This structural analysis indicates that the PB2 domain forms with PB1 an extended basic cluster at the NC interface of septin 9.

### Contribution of the PB domains to septin 9 membrane association

To determine whether the PB domains are equally involved in septin 9-membrane interaction, we first did MD simulations of the interaction of the monomeric structure of the protein shown in Figure 1D with membranes devoid of PtdIns4P (Figure S2A). The simulations were done by placing the protein close to a dioleoylphosphatidylcholine (DOPC) membrane and leaving it with the possibility to change conformation over time. We simulated three conditions by changing the initial protein conformation and velocity (Figure S2A, MD1-3). In all cases, septin 9 was recruited to the membrane. However, we found that PB1 was always in contact with the membrane (Figure S2A)(see experimental procedure), while PB2 was not in one of the conditions (Figure S2A, MD1). This observation suggests that PB1 may be better positioned for interacting with membranes.

To further distinguish the implication of PB1 and PB2 in septin 9 interaction with PI lipids, we took advantage of the recently resolved crystal structure of septin 3 (Macedo et al., 2013), which belongs to the same subgroup as septin 9 (Figure 1A). By using homology modeling, we built a septin 9 monomer and tetramer (Figure S2B,C). We used the tetramer to determine its spatial organization on a DOPC/DOPE bilayer containing PtdIns5P (Lomize et al., 2012). In the membrane-proximal PB1-PB2 cluster of the tetramer, we found that the basic residues R^289^, R^290^, K^291^ and K^400^ were particularly well positioned to interact with the phosphate headgroup of the phosphoinositides (Figure 2A). Three of these residues, namely R289, R290, and K291, belong to PB1, and K400 to PB2. This analysis also supports that PB1 is more involved in regulating septin 9 membrane binding than PB2, consistent with the previous MD results (Figure S2A).

**Figure 2:**
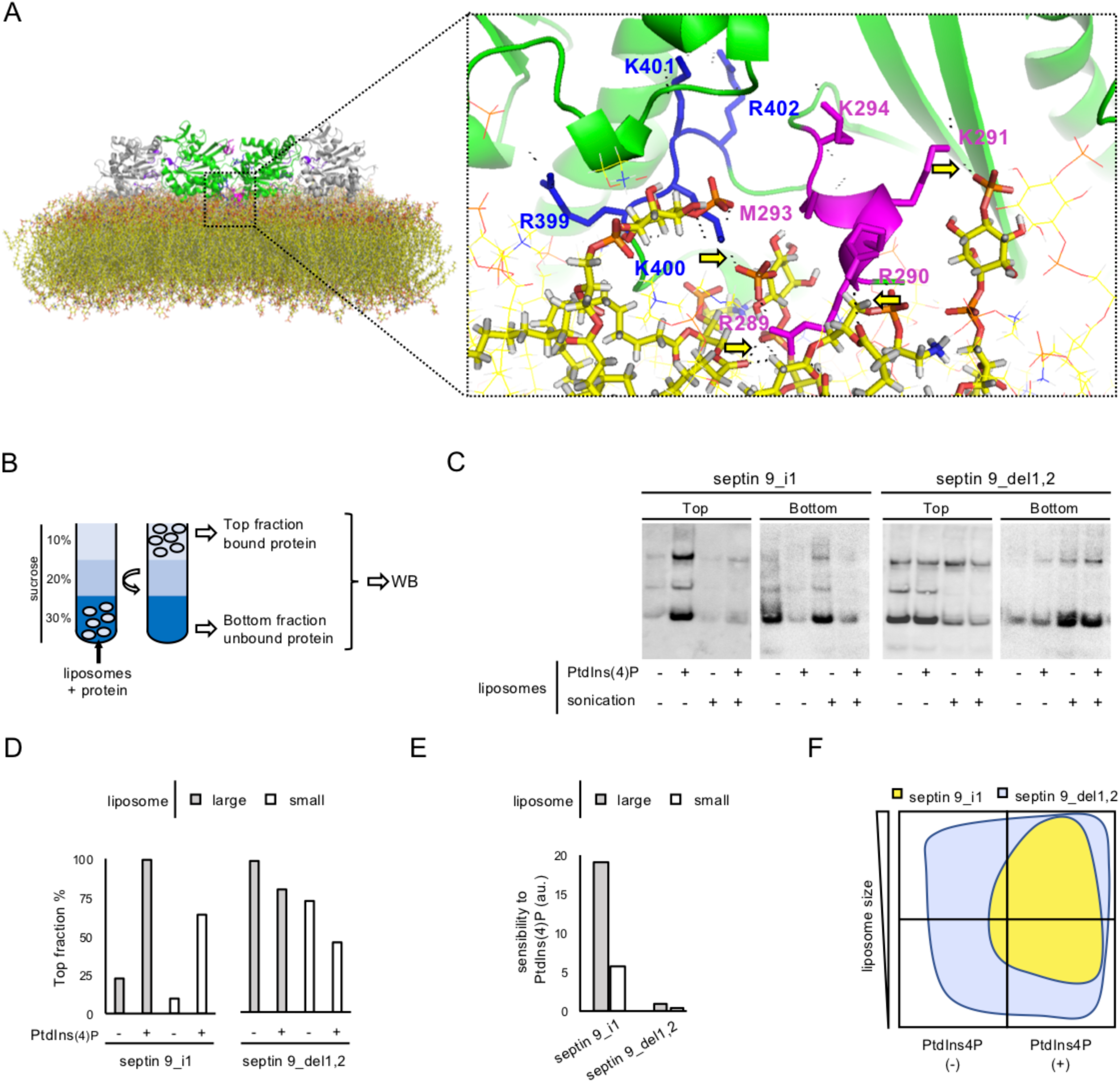
PBs are required for septin 9_i1 PIs specific interaction and membranes form recognition. A. Modeling of septin 9 complex interaction with a patch of PI containing lipid bilayer. The dotted rectangle indicates the PB1-PB2 area proximal to the membrane shown in higher magnification. The residues for PB1 and PB2 are labeled. Yellow arrows indicate the interaction between PBs and the membrane. B. Schematic representation illustrating the liposome floatation assay C. Western blots of septin 9_i1 (I1) and septin 9_del1.2 (Del1,2) subjected to a liposome flotation assay; the arrow indicates the band corresponding to Septin9_i1 V5 tagged (68 kDa) further analyzed. D. Bar graph representing the percentage of protein in the top fraction (bound protein) from the analysis of the blots presented in panel B. E. Bar graph representing the sensitivity of the protein to PtdIns4 (the ratio of protein bound with PtdIns4P(+) liposomes to protein bound with PtdIns4P(-) liposomes). Dashed line indicates the 1 value. F. Schematic phase diagram illustrating the result of Western blot analysis of the liposome flotation assay.

### Septin 9 interaction with membranes *in vitro* does not require any of the PB domains

We decided to study the contribution of the PB domains in septin association *in vitro*. Big and small liposomes (e.g. above 300 nm and below 200 nm) containing or not PtdIns4P were respectively generated by vortex and high power sonication, to respectively mimic flat and curved membranes. The liposomes were then mixed with septin 9_i1, and septin 9_del1,2. A sucrose gradient flotation assay was subsequently performed (Figure 2B). In this assay, only bound proteins float up with the liposomes to the top fraction (Figure 2B). The supernatant fraction was collected and the amount of bound proteins determined by Western blotting (Figure 2C).

As expected, septin_i1 bound stronger to liposomes containing PtdIns4P than to those lacking PtdIns4P (Figure 2C-F). However, binding was reduced on smaller liposomes (Figure 2C-F, Figure S2D), which suggests that septin 9 had more binding capacity to flat than to positively curved membrane regions. Surprisingly, we found that septin 9_del1,2, which lacked both PB domains, efficiently bound to all membranes (Figure 2C-F), despite that it also had a slight preference to flatter membranes (Figure S2D). This efficient binding of septin 9_del1,2 was not detected in the PIP strip overlay assay because perfect membrane bilayers are not generated in this method (Figure 1C); it was however consistent with the binding of septin 9 to membranes devoid of PI lipids by MD simulations (Figure S2A). Finally, the binding of septin 9_del1 and septin 9_del2 was not enhanced by the presence PtdIns4P, and it was almost lost on small liposomes (Figure S2E). These results suggest that PB1 and PB2 are both required for the specific and efficient binding of septin 9 to PI-containing membranes.

In conclusion, our above data, and particularly those obtained with septin 9_del1,2, suggest that septin 9 can bind bilayer membranes without involving its PB domains. Both PB domains seem to be necessary to provide septin 9 binding selectivity to PI-containing membranes, and especially to flat membranes or at least to non-positively curved ones.

### Septins have PB-adjacent amphipathic helices (AH)

We next asked how Since septin 9 was able to interact directly with membranes in the absence of PI lipids and without its PB domains (Figure 2). Many soluble proteins bind membranes using amphipathic α-helix motifs (AHs), which are moreover able to detect subtle differences in membrane curvature and charges (Bigay and Antonny, 2012), as septin 9 did. We therefore did a bioinformatics screening for AHs in the full sequence of septin 9. Two striking sequences emerged from our analysis as being the most prominent AHs (aa274 to 294, aa370 to 402). Surprisingly, these AHs were directly adjacent to PB1 and PB2 respectively (Figure 3A). The sequence close to PB1 corresponds to a flexible strand that can fold into an AH, which is probably why it is missing in the septin 9 crystal structure. In MD simulation, we found that this portion of the protein indeed folded as an α-helix (Figure S1D) well positioned to bind membranes (Figure 3A). The AH close to PB2 is folded but oriented toward the interior of the protein, unless a conformational switch happens. These AHs contain very hydrophobic residues, such as tyrosine, tryptophan and phenylalanine, which is a feature enhancing membrane association. Thus, our analysis suggests that septin 9 has at least one PB-adjacent AH, associated to PB1, which can mediate its physical association with membranes. In other septins, we also found flagrant AHs juxtaposed with the PB2 of septin 2, 6, and 7 with which septin 9 interacts to form the octamer. We took advantage of the existence of a crystal structure of the trimer formed by septin 2-6-7 to verify the orientation of the AHs. We found that the AH-PB2 of septin 6 was suitably oriented to bind membranes (Figure 3B). The AH feature at the PB1 domain of these septin 9 counterparts were less pronouced (Figure 3B, Figure S3A-D).

**Figure 3:**
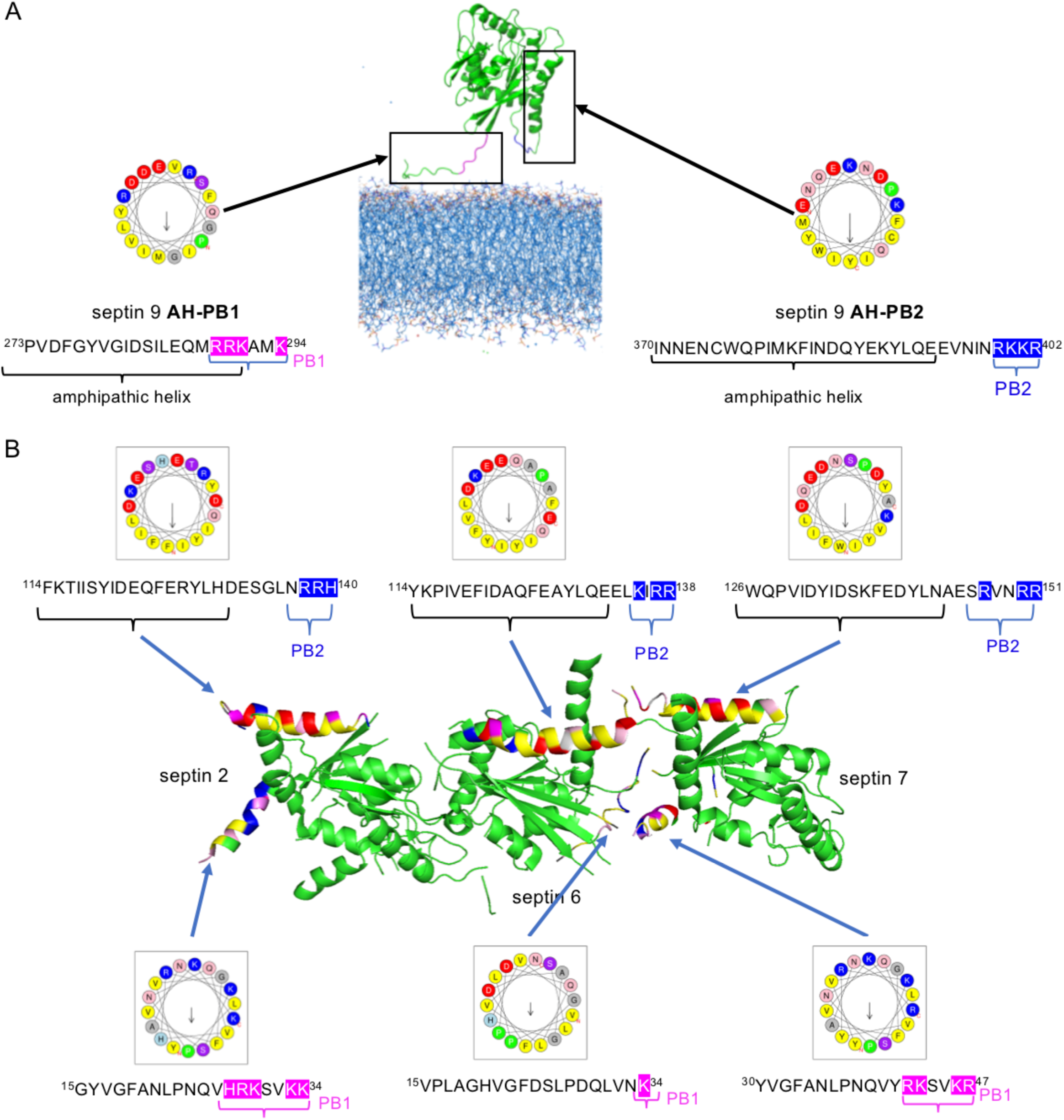
Septin 9 has putative PB-associated amphipathic helices mediating its binding to the membrane. **A.** Crystal structural model of septin 9 (PDB code 5cyp) showing the predicted amphipathic helices opposed to PB1 and PB2 and their helical wheel representation generated using Heliquest. **B.** Structural model of the (NC2G/G6NC/NC7G) septin complex published by Sirajuddin et al. showing the predicted amphipathic helices opposed to PB2 and the helical wheel representation of these helices generated using Heliquest. The sequence of each predicted helix is presented below, the corresponding helical wheel and PB1 basic residues are highlighted in magenta and those of PB2 in blue.

### Septin 9 PB domains are essential for the formation of septin filaments

Our data supports that the AHs of septin 9 are probably the membrane binding motifs which are restricted to PI-containing membranes by the PB domains. In this scenario, the PB domains would be more available to participate in the NC/NC interactions mediating the formation of septin high order structures.

We transfected HeLa cells with septin 9_i1, which we found incorporated in high order filamentous structures that also contained endogenous septin 9 and septin 2 (Figure 4A, B). However, the high expression of septin 9_i1 seemed to displace endogenous septin 9 from the filaments, while septin 2 remained recruited at similar level (Figure 4B,C). Our overexpressed septin 9 construct thus had a dominant negative effect on endogenous septin 9 (Figure S4E). When cells were transfected with the PB-deleted constructs (Del1, Del2, Del1,2), the filamentous structures were lost (Figure 4D, E, F) despite the presence of the endogenous septin 9. Here also, these constructs have the dominant negative effect of septin 9 (Figure S4E), and septin 2 was not present in filaments (Figure 4D). These results suggest that both PB domains are involved in septin 9 assembly, in line with our previous results from the structural analysis (Figure 1E, Figure S2C).

**Figure 4:**
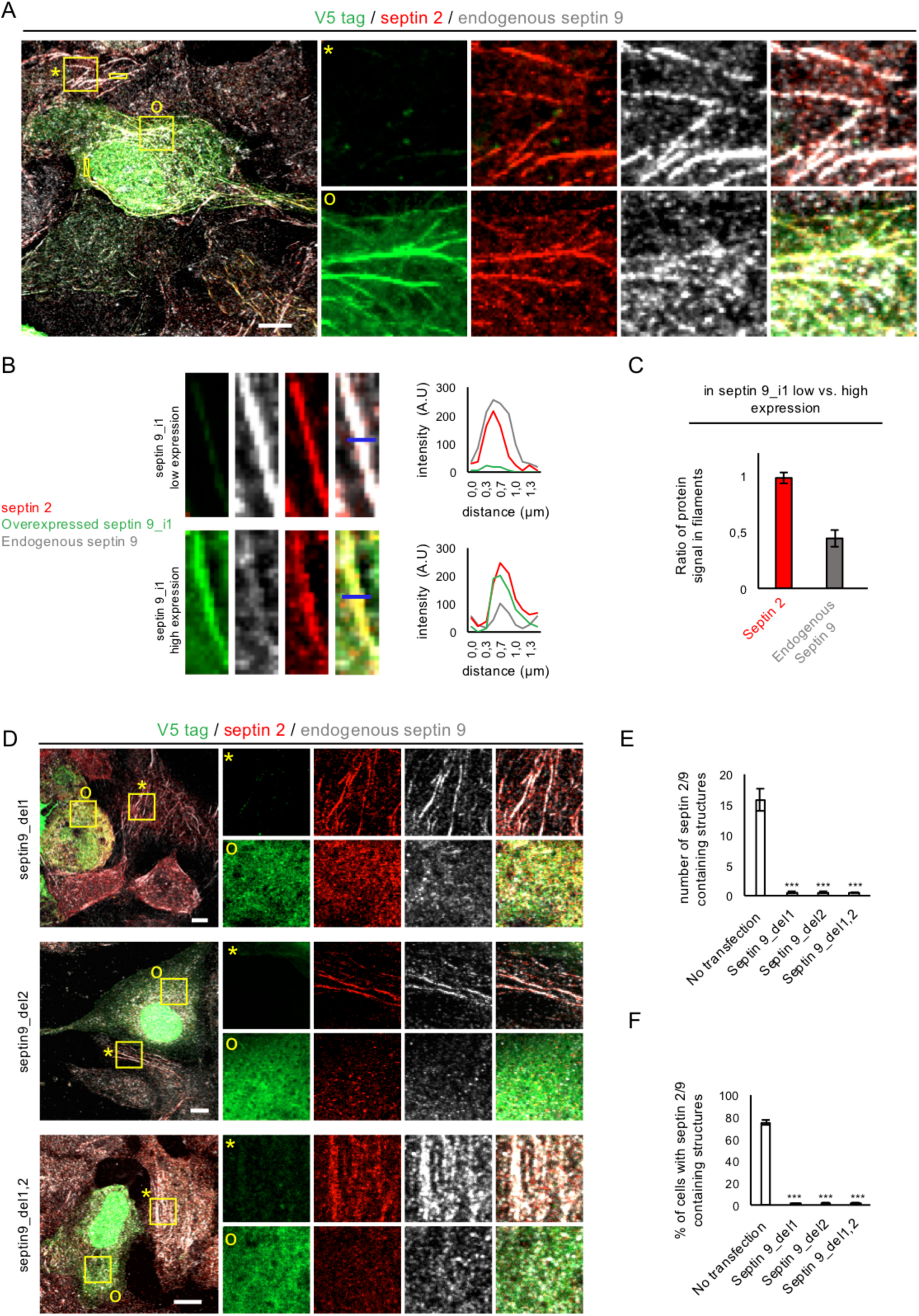
Overexpressed septin 9_i1 replaces indigenous septin 9 in septin filament while PBs muted septin 9 expression impairs them. A. Huh7,5 cells transfected with septin9_i1 for 48h then fixed and stained for V5 tag in green (septin 9_i1) endogenous septin 2 (red) and endogenous septin 9 (grey). (*) indicates septin9_i1 low expression cell and (0) indicates septin9_i1 high expression cell. Squares indicate the area shown at higher magnification on the right. Scale bar: 10µm. B. The small yellow rectangles shown in panel A are shown here in higher magnification. On the right, the line graphs show the line profile analysis of the lines shown in these images. C. Bar graph showing the ratio of the intensity of endogenous septin 2 and endogenous septin 9 in septin 9_i1 low expression cells to those of septin 9_i1 high expression. Values are mean ± SEM from 10 filament under each condition form tow independent experiments. D. Huh7,5 cells transfected with septin 9_del1 (Del1), septin 9_del2 (Del2) or septin 9_del1,2 (Del1,2) for 48h, then fixed and stained for V5 tag in green, endogenous septin 2 (red) and endogenous septin 9 (grey). (*) indicates a low expression or not transfected cell and (0) indicates a transfected cell. Squares indicate the area shown at higher magnification to the right. E. Bar graph representing the number of the filament structures containing both endogenous septin 9 and septin 2. Values are mean ± SEM from 10 cells under each condition form tow independent experiments. F. Bar graph representing the percentage of cells containing filament structures of endogenous septin 9 and septin 2. Values are mean ± SEM from 50 cells under each condition form tow independent experiments.

The avoid possible conformational changes of the protein and its dysfunction due to the deletion of the PB domains, we did simple point mutations. We substituted the basic lysine and arginine amino acids (K and R) by glutamine (Q), which is a non-charged but polar amino acid, with a long side chain as in K and R; this substitution is therefore optimal for minimizing possible conformational changes in the protein. For example, in septin 9_Q1, residues R and K of PB1 were not deleted as in septin 9_del1 but replaced by Q. We observed that cells transfected with septin 9_Q1, septin 9_Q2, or septin 9_Q1,2 were unable to form septin filaments (Figure S4A-C), as with the deletion (Figure 4D-F). We next did simple mutations by substituting one or two R with alanine (A) within PB1 and also lost the filaments (Figure S4D). All these constructs did not affect the normal expression of endogenous septin 9 (Figure S4E).

In conclusion, the point mutant phenocopied the deletion constructs and the results suggest that the positively charged PB cluster at the NC/NC interface is essential to the formation of septin filaments.

### Septin 9 PB domains are critical for Golgi assembly

Septin 9 binds mainly to PtdIns3P, PtdIns4P and PtdIns5P which are respectively detected primarily in the early endosomes (EE), the Golgi apparatus and the endoplasmic reticulum (ER) (Pendaries et al., 2006; Kutateladze, 2010; Sarkes and Rameh, 2010). We thus wanted to probe whether PB1 and PB2 contribute to the organization of these endomembrane compartments. We expressed septin 9_i1 and the mutant constructs in HeLa cells and studied the morphology of these organelles.

A significant increase of EE and ER markers was observed in the perinuclear region of septin 9_i1-expressing cells when compared to EV and septin 9 mutants, which were similar (Figure S5A-E). The most striking observation of the deletion of the PB domains was on the Golgi structure (Figure 5A, B). In cells expressing septin 9_i1, we found the normal phenotype of a compact Golgi embedded in high order structures of septin 9_i1 filaments (Figure 5A, C), which contained septin 2, 6 and 7 (Figure S5F). Strikingly, the deletion or mutation of any of the PB domains led to Golgi fragmentation (Figure 5A, B, Figure S5G).

**Figure 5:**
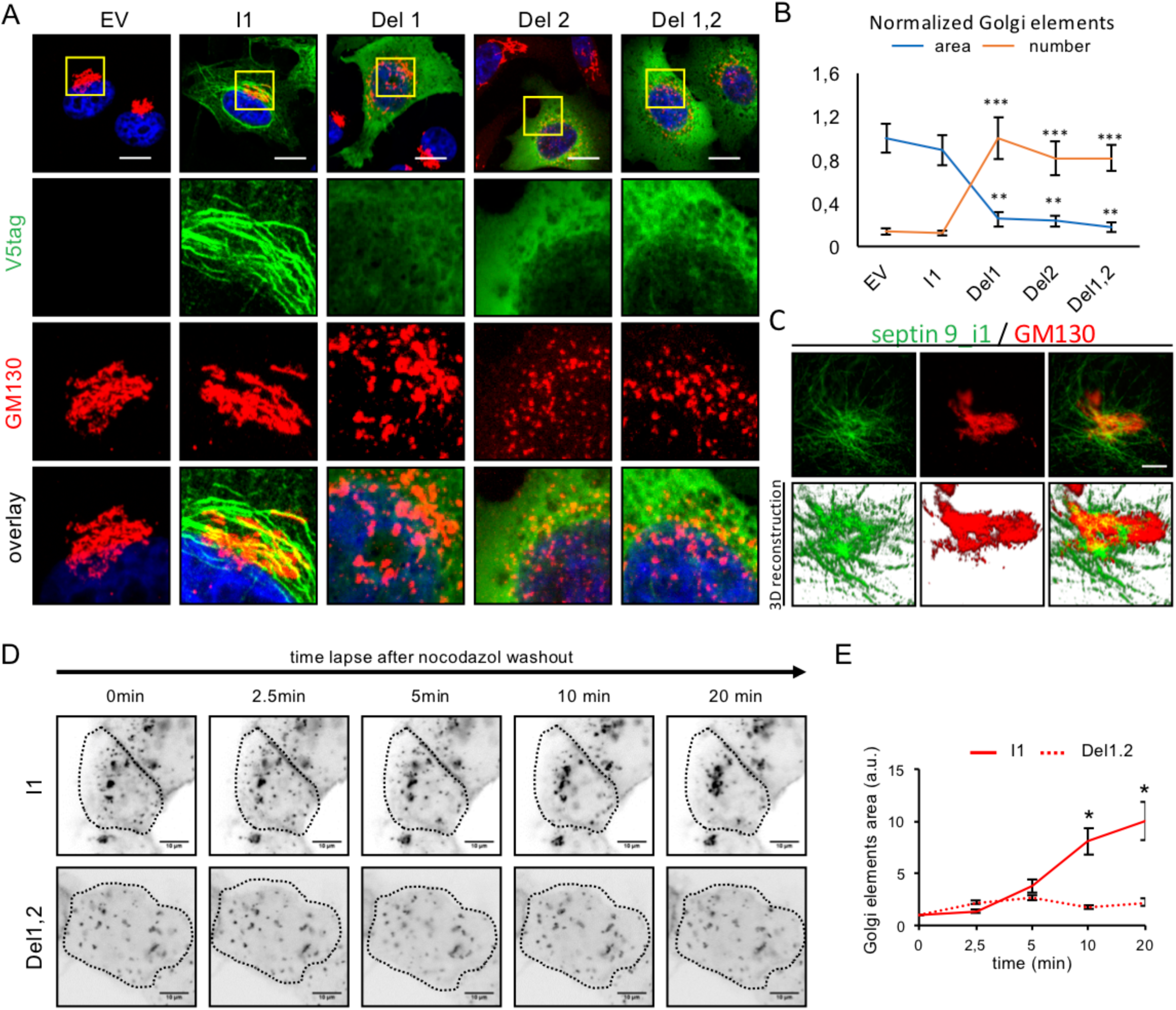
Septin 9 localizes to Golgi and is required for its compact morphology. A. HeLa cells transfected with either an empty vector (EV), septin 9_i1 (I1), septin 9_del1 (Del1), septin 9_del2 (Del2) or septin 9_del1,2 (Del1,2) for 48h, then fixed and stained for Golgi with GM130 (red) and V5tag (green). Squares indicate the area shown at higher magnification below. Scale bar: 10µm. B. Line graph representing normalized number and size of Golgi elements. Values are mean ± SEM from 15 cells under each condition for three independent experiments. **P<0.001, ***P<0.0001, (Student’s t-test). C. STED high resolution microscopy images of HeLa cells transfected with septin 9_i1 (I1) then fixed and stained for GM130 (red) and V5tag (green). Images were shown in 3D reconstruction images below. Scale bar: 5µm. D. MDCK cells stably expressing septin 9_i1 or septin 9_del1,2 and transfected with KDE-GFP for 24h subjected to nocodazole washout. Images represent video frames illustrating the reassembly of Golgi after the removal of nocodazole. E. Dotted lines indicate the cell periphery. Line graphs to the right represent fold-increases in Golgi elements relative to time 0 during three experiments. Scale bar: 10µm. *P<0.05, (Student’s t-test).

Assembly of the Golgi apparatus is dependent on microtubule polymerization (Miller et al., 2009). The depolymerization of microtubules, typically under nocodazole treatment, results in Golgi fragmentation; removal of the nocodazole enables the re-polymerization of the microtubules and Golgi reassembly. We thus took MDCK cells stably expressing septin 9_i1 and septin 9_del1,2 and treated them with nocodazole to induce Golgi fragmentation. The nocodazole was then washed out and the Golgi reassembly monitored (Figure 5D). In septin 9_i1 cells, the scattered Golgi elements reassembled normally within 20 min (Figure 5D, E) but not in septin 9_del1,2 cells (Figure 5D, E), even at much longer time points (Figure S5H).

These above observations suggest a function of septin filaments, which depends on septin 9 PB domains, on Golgi assembly.

### Mutations of the PB domains cause Golgi fragmentation but not Septin 9 dissociation from membrane

Our data suggests that the lack of PB domains should induce the loss of specific binding of septin 9 to PtdIns4P (Figure 2). We decided to follow in cells the interaction of septin 9 with PtdIns4P. HeLa cells were transfected with cDNAs of EV, septin 9_i1 and Del1, Del2 or Del1,2, and stained for V5 tag, TGN46 (Golgi marker) and PtdIns4P (Figure 6A and Figure S6A). PtdIns4P was strongly colocalized with compact Golgi in EV and septin 9_i1, as expected (Dippold et al., 2009). In cells transfected with Del1, Del2 or Del1,2 constructs (Figure 6A), the Golgi was fragmented, as seen previously, but no significant difference in co-localization between TGN and PtdIns4P was noticed (Figure 6B). However, we observed a non-significant decrease in the co-localization of septin 9 with PtdIns4P (Figure 6C) between the various transfected constructs. The PB-deleted mutants still displayed important signals on small spherical compartments which were possibly Golgi ministacks, vesicles or other structures (Figure 6A, Figure S6A). To determine whether the mutant septin 9 proteins were associated to these membrane structures, we permeabilized the cells and removed the soluble proteins prior to fixation. We observed that the septin 9 mutated protein signal remained onto intracellular structures (Figure 6D). This observation suggests that the lack of PB domains does not prevent the binding of the mutant proteins to possible membrane structures, in line with our in vitro assays which revealed the ability of the mutants to bind to membranes (Figure 2). To further test this finding, a subcellular fractionation assay was done (Figure S6B). We found that septin 9_i1 always peaked with the Golgi marker, while septin 9_del1,2 had a more spread out signal, suggesting that it probably bound to other membranes, including fragmented Golgi elements (Figure S6B).

**Figure 6:**
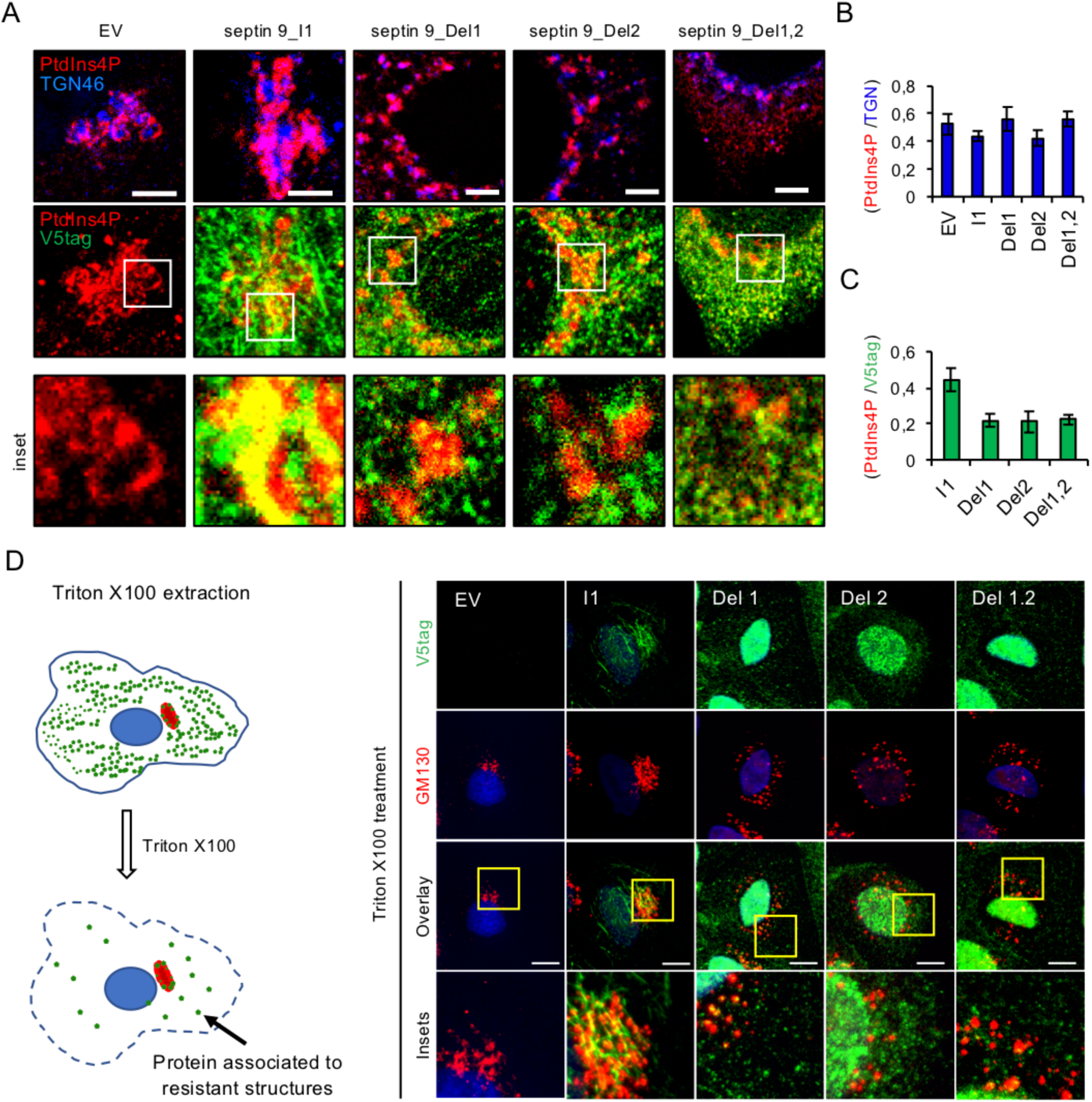
PBs are required for the specific recruitment of septin 9_i1 to the Golgi. A. HeLa cells were transfected with either an empty vector (EV), septin 9_i1 (I1), septin 9_del1 (Del1), septin 9_del2 (Del2) or septin 9_del1,2 (Del1,2) for 48h before being fixed and stained for PtdIns4P (red), TGN46 (blue) and V5tag (green). Squares indicate the area shown below at higher magnification. Scale bar: 3µm. B. Bar graph represents Pearson’s correlation coefficient (Rr) analysis of PtdIns4P and TGN46. Values are mean ± SEM of 10 cells under each condition from three experiments. C. Bar graph representing Pearson’s correlation coefficient (Rr) analysis of PtdIns4P and V5 tag. Values are mean ± SEM of 10 cells under each condition from three experiments. D. Left, schematic representation illustrating cell extraction with triton X100. On the right, MDCK stably transfected with either EV, septin 9_i1 (I1) or septin_9 del1 (Del1) septin_9 del2 (Del2), septin_9 del1.2 (Del1,2) were grown for 24h on coverslips. The cells were then extracted using cold PBS buffer containing triton X100 at 0.1% for 30 seconds then fixed and stained for V5tag (green) and GM130 (red). Scale bar: 10µm.

Taken together, our results suggest that PB1 and PB2 restrict septin 9 binding to PI-containing membranes, such as PtdIns4P on Golgi, but are not directly responsible for septin membrane association. They have a major function in septin complex assembly and subsequent organelle structuration.

### Endogenous septin 9 localizes to Golgi and regulates its compactness and functionality

The transfection of our septin 9 mutants constructs, which had a dominant negative effect on endogenous septin 9, could be the reason of the Golgi fragmentation. We thus studies the endogenous septin 9 and found that it also colocalized with Golgi (Figure 7A). This observation was likewise supported by a subcellular fractionation assay (Figure 7B) in which both septin 2 and septin 9 peaked with GM130 (Figure 7B). These results suggested that endogenous septin 9 is present in septin structures associated with the Golgi.

**Figure 7:**
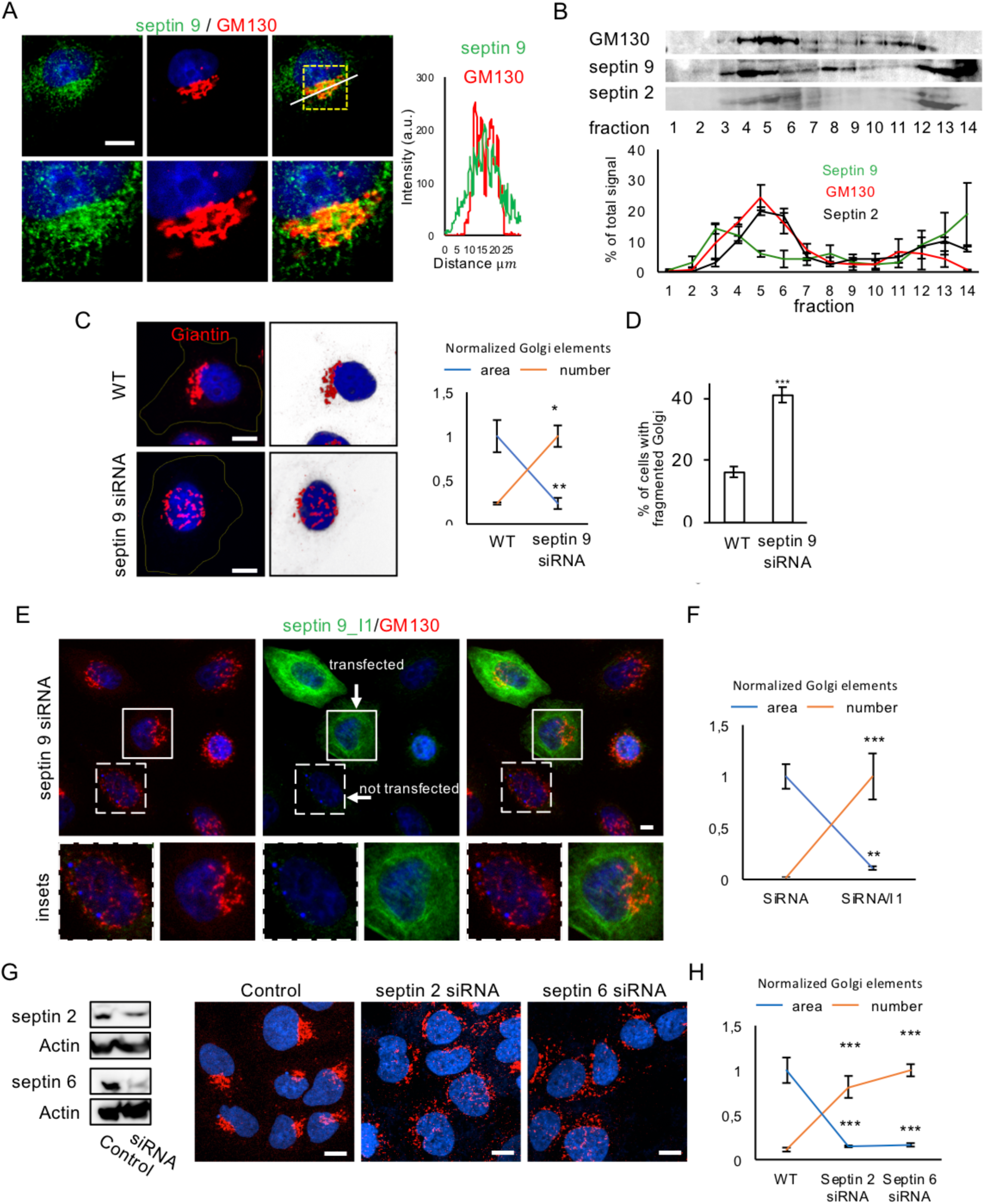
Septin 9 localizes to Golgi and is required similarly to septins for its compact morphology. A. Huh7 cells Fixed and stained for septin 9 and GM130. Scale bar: 5µm. On the right, line profile of the white line shown in the merge image. B. Huh7 cells were grown for 48h before being subjected to a subcellular fractionation assay and analyzed with western blot for septin 9 and GM130 and septin 2. Line graph below shows the densitometry analysis of subcellular fractionation assay. Values are mean ± SEM from three independent experiments. C. Septin 9 siRNA and control cells stained for giantin (red). Cells are shown in 2D images with a black background and in 3D reconstruction images with a white background. Scale bar: 10µm. On the right line graph represents normalized size and number of Golgi elements. Values are mean ± SEM of 150 cells from three independent experiments. *P<0.05, **P<0.001, (Student’s t-test). D. Percentage of cells with fragmented Golgi obtained from three independent experiments. At least 800 cells were counted for each condition. Values are mean ± SEM. ***P<0.0001, (Student’s t-test). E. septin 9 siRNA cells were transfected with septin 9_i1 and stained for GM130 (red) V5tag (green). Squares indicate the area shown at higher magnification below. Scale bar: 10µm. F. Bar graph representing the size and number of Golgi elements in 20 cells from two independent experiments done as described in E. **P<0.001, ***P<0.0001, (Student’s t-test). G. Huh7.5 cells transfected with septin 2 siRNA or septin 6 siRNA for 48h were analyzed with western blot (left) and confocal microscopy for GM130 (red) (right). Scale bar: 10µm. H. Line graph representing normalized size and number of Golgi elements in 10 cells from two independent experiments. ***P<0.0001, (Student’s t-test).

We next worked with a cell line stably transfected with septin 9 siRNA (Figure 7C). In control cells, Giantin, which is also a Golgi marker, had a polarized and compact feature, i.e. enriched on one side of the nucleus (Figure 7C). Instead, in the septin 9 siRNA cells, we observed a fragmentation of the Golgi (Figure 7C). Interestingly, this fragmentation was rescued by transfection with septin 9_i1 (Figure 7 E, F), but not with the PB-deleted constructs (Figure S7A, B). Here also, we performed the nocodazole washout assay which confirmed that the Golgi reassembly is dependent on endogenous septin 9 (Figure S7C). Thus, septin 9 has an important role in maintaining Golgi compact structure.

Our overall data suggest that septin 9 (endogenous and septin 9_i1) is incorporated into filaments which are critical to Golgi structure. Since septin 2 and Septin 6 are involved in these filaments, they should also control Golgi compactness. We accordingly found that their depletion by a specific siRNA promoted Golgi fragmentation (Figure 7G, H, Figure S7E). This data reinforced our model in which the septin complexes are incorporated into filament structures which are associated with the Golgi membrane and control its compactness.

Finally, fragmentation of the Golgi apparatus is known to impede on the secretory pathways (Xiang et al., 2013; Lavieu et al., 2014; Joshi et al., 2015). We thus probed whether septin 9 depletion (Figure S8A,B) impeded secretory pathways. We first used a Venus-tagged Neuropeptide Y construct (NPY-venus) as a model of a secreted protein; we determined its secreted level in the cell culture medium and lysates. We found that septin 9 depletion resulted in a significant decrease of secreted NPY-Venus (Figure S8C). We next studied the trafficking of a temperature-sensitive variant of the vesicular stomatitis virus G protein tagged with GFP (tsVSVG-GFP) (this protein remains in the ER at 40°C but enters the secretory pathway when the temperature is lowered to 32°C (Presley et al., 1997)). Control and septin 9 depleted cells were transfected with tsVSVG-GFP and fixed at different time points after temperature was lowered from 40°C to 32°C (Figure S8D). After 2 hours, the protein signal in the Golgi area was greatly reduced in control cells whereas it remained intact in septin 9 depleted cells, suggesting a defect in intracellular trafficking. Taken together, these data indicated that septin 9 plays a critical role in Golgi compactness and subsequent function in cellular trafficking.

## Discussion

Septins belong to the family of GTPase proteins which assemble into macrostructure filaments which are important for membrane remodeling processes. Here we have shown that septin 9 is particularly necessary for Golgi structure and function, as the absence of septin 9 provokes Golgi dissociation and impairs secretory pathways. Our results support the idea that septin 9 and other septins, such as septin 2 and septin 6, form a filament matrix which harbors Golgi stacks.

The association of septins with membranes was previously thought to be specifically mediated by an interaction of their polybasic domain PB1 with PI lipids (Pan et al., 2007; Zhang et al., 1999; Tanaka-Takiguchi et al., 2009; Casamayor and Snyder, 2003). Here we found that septin membrane association is much more subtly regulated. First, we identified the presence of a second polybasic domain (PB2) in septin 9, which is conserved among the different human septins and regulates the interaction of septin with PI lipids. Second, we identified AH structures adjacent to PB1 and PB2 which probably mediate the physical association of septin 9 with membranes. Our result support that PB1 and PB2 would act together to restrict the binding of these AHs of septin 9 to PI-containing membranes, particularly to non-positively curved membranes. This conclusion is in line with the accumulation of septins *in vivo* to specific plasma membrane ingressions or concavities, such as cleavage furrows (Spiliotis and Nelson, 2006). Finally, based on our *in vitro* observations, MD (Figure S2A) and the modeling of the interaction of septin 9 complex with PtdIns(4)P containing membrane (Figure 2A), PB1 appears to be more closely involved in regulating septin 9 membrane binding specificity than PB2, but both are critical for septin filament assembly.

Whether in a cellular context septin 9 acts in a monomeric form is not known. Except for the septin 9 isoform septin 9_i4, which have been found in non-filament structures (Chacko et al., 2005), most septins have been so far reported incorporated into filamentous structures, and whether they exist under monomeric form is also unknow. Despite that our *in vitro* studies were done with the monomeric septin 9, our results give information on how septin 9 membrane binding determinants might influence the localization of septin oligomers to membranes. These results are consistent with a recent report of the presence of AHs structures in oligomeric septin filaments able to sense macroscopic curvatures (Cannon et al., 2018).

Based on our results, we propose that septin 9 binds membranes with AHs that can strongly associate to the membrane. In this setting, PB domains may interact with PI lipids to dock the protein on specific organelles, thereby preventing non-specific binding. This mechanism would enrich the protein onto the membrane, thanks to the PBs, and stabilize it thanks to the AHs.

It is not clear yet how septins control the Golgi structure. An hypothesis is that the PB domains contribute to the enrich septin 9 or the octameric complexes to Golgi elements. Assembly of the octamers would form the septin filaments, which will in the meantime bring different Golgi elements closer enough to promote their fusion, e.g. by SNARE proteins. In this view, a septin matrix will form a structure embedding the Golgi elements. This model is consistent with our experiments on Golgi reassembly after nocodazole treatment: reassembly occurred with septin 9_i1, whereas with Del1,2 (Figure 5D, E, Figure S5H) or septin 9 depletion (Figure S7C) no reassembly occurred. Consistent also with our model, the Golgi is fragmented by the depletion of septin 2 or 6. Hence, our study reveals the importance of the high order septin structures in Golgi homeostasis. Since microtubules and actin filaments are also known to maintain Golgi structure and interact also with septins (Miller et al., 2009; Egea et al., 2006; Kondylis et al., 2007; Fung et al., 2014), it is plausible that septins are engaged into hybrid filaments with microtubules and actin for maintaining Golgi structure.

On other organelles, we observed that PB1 and PB2 affected ER and EE organization, which was reminiscent to the role that we described for septin 9 in lipid droplet dynamics; interestingly, septin filaments were prominent around large lipid droplets (Akil et al., 2016). This particular localization of septin high order structures to sites of micron-scale membrane curvature is emerging as an important feature of organelle dynamics (Cannon et al., 2017).

Finally, septin structures have been proposed to act as a physical barrier for the non-specific docking of vesicles to the active zone of the synapse (Yang et al., 2010). Thus, septin structures could also behave as physical barriers that prevent the collapse of the Golgi stacks or their connection to other organelles. Our findings on the role of septin 9 on Golgi could probably be extended to other organelles and offer a new paradigm to the structural and biological functions of septins.

## Author contributions

A.G.D., M.O. and A.R.T. designed the research, analyzed the data and wrote the paper. M.O. conducted the experiments, with the support of C.P, B.G., N.B., D.S., C.T. and A.R.T., C.T. purified the recombinant proteins. A.S.C. and RCG analyzed septin 9 crystal structures and performed the molecular dynamics simulations using these structures. C.T and S.B with the help of R.G., J.Y. and T.T. performed the septin 9 homology modeling and septin 9/PIs interaction modeling.

### Acknowledgements

This work received support in the form of a grant from the Association pour la Recherche sur le Cancer (ARC/SUBV/CKLQ6) for AGD, and from the ANRS (France Recherche Nord & sud SIDA – hiv Hépatites: FRENSH) through a grant to SB and a PhD fellowship for CT. A.R.T is supported by the ANR-TERC (LDEN), Paris Sciences et Lettres, and ATIP-Avenir. We would like to thank H. Russell (Queen’s University, Belfast, UK.) for providing the septin 9_i1 construct. We also thank A. Baillet (Université Paris-Sud, UMR-S 1193, Villejuif, France) for access to live cell imaging facilities.

