## Supplementary material for "Two polybasic domains of septin9 control septin filaments assembly and Golgi integrity": Supplimental Informations (from another version of this preprint: version 1, 2019-01-30)

### Supplemental Information

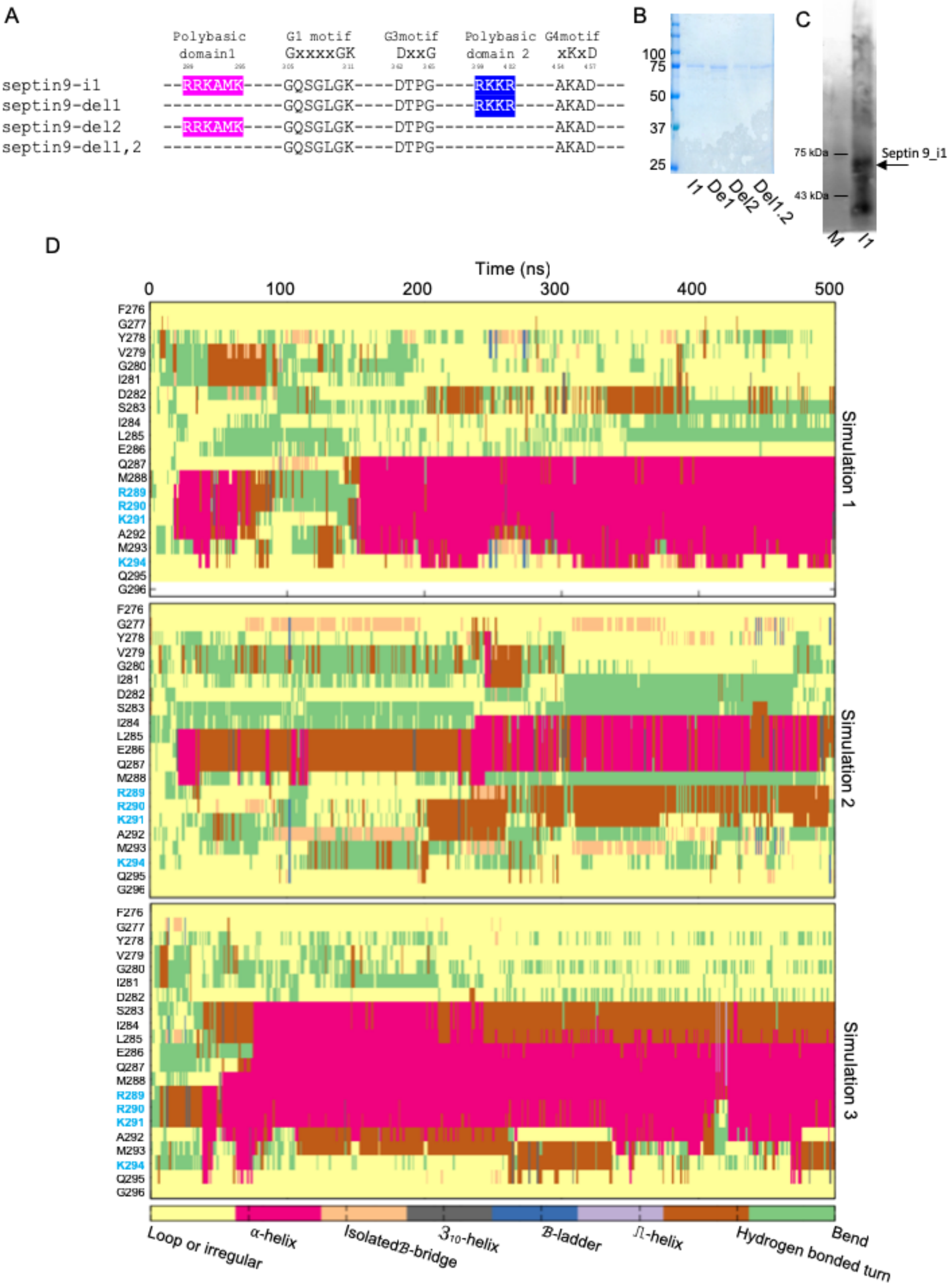


Supplementary Figure **(Related to Figure 1)**

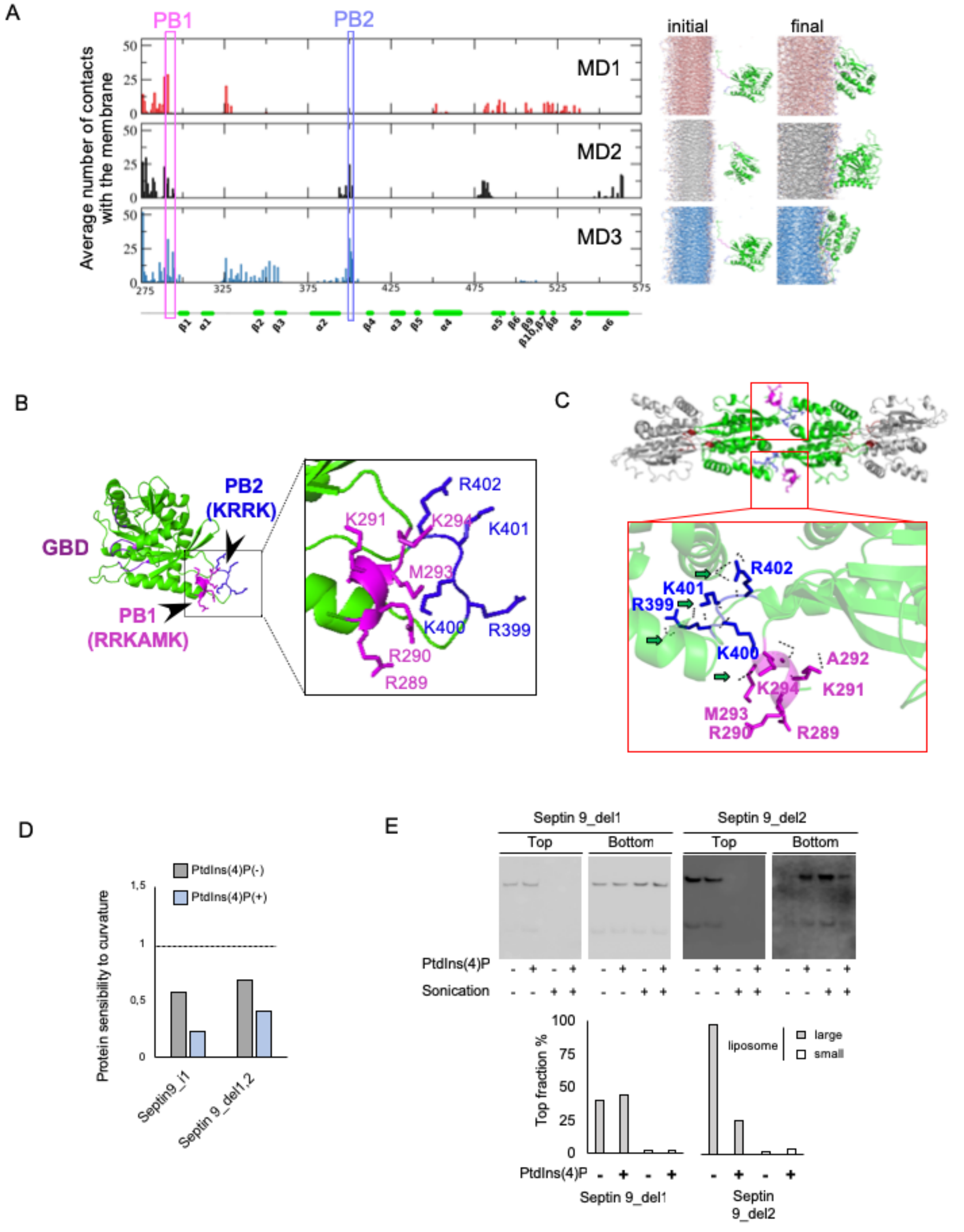


Supplementary Figure 2 **(Related to Figure 2)**

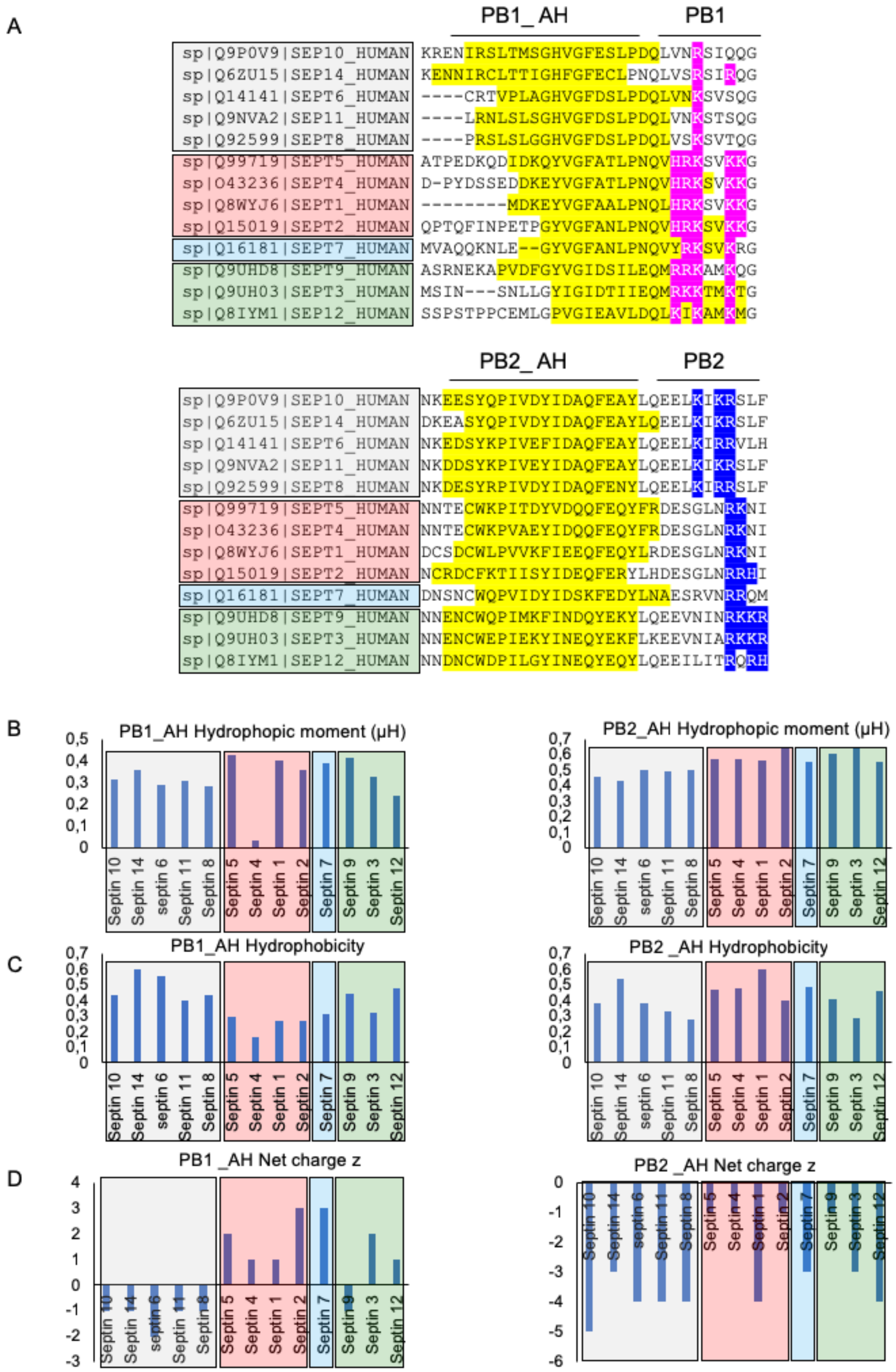


Supplementary Figure 3 **(Related to Figure 3)**

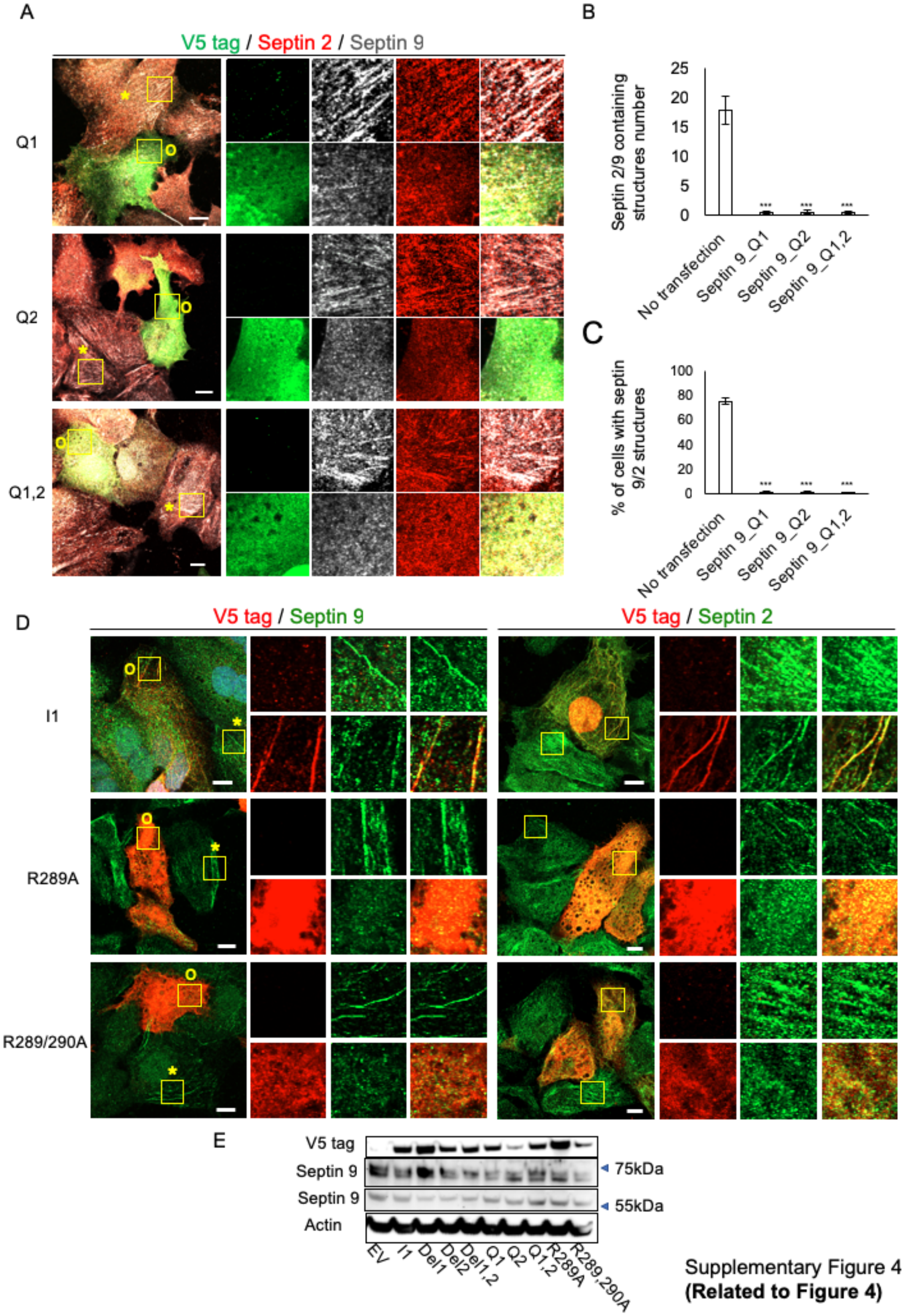


Supplementary Figure 4
**(Related to Figure 4)**

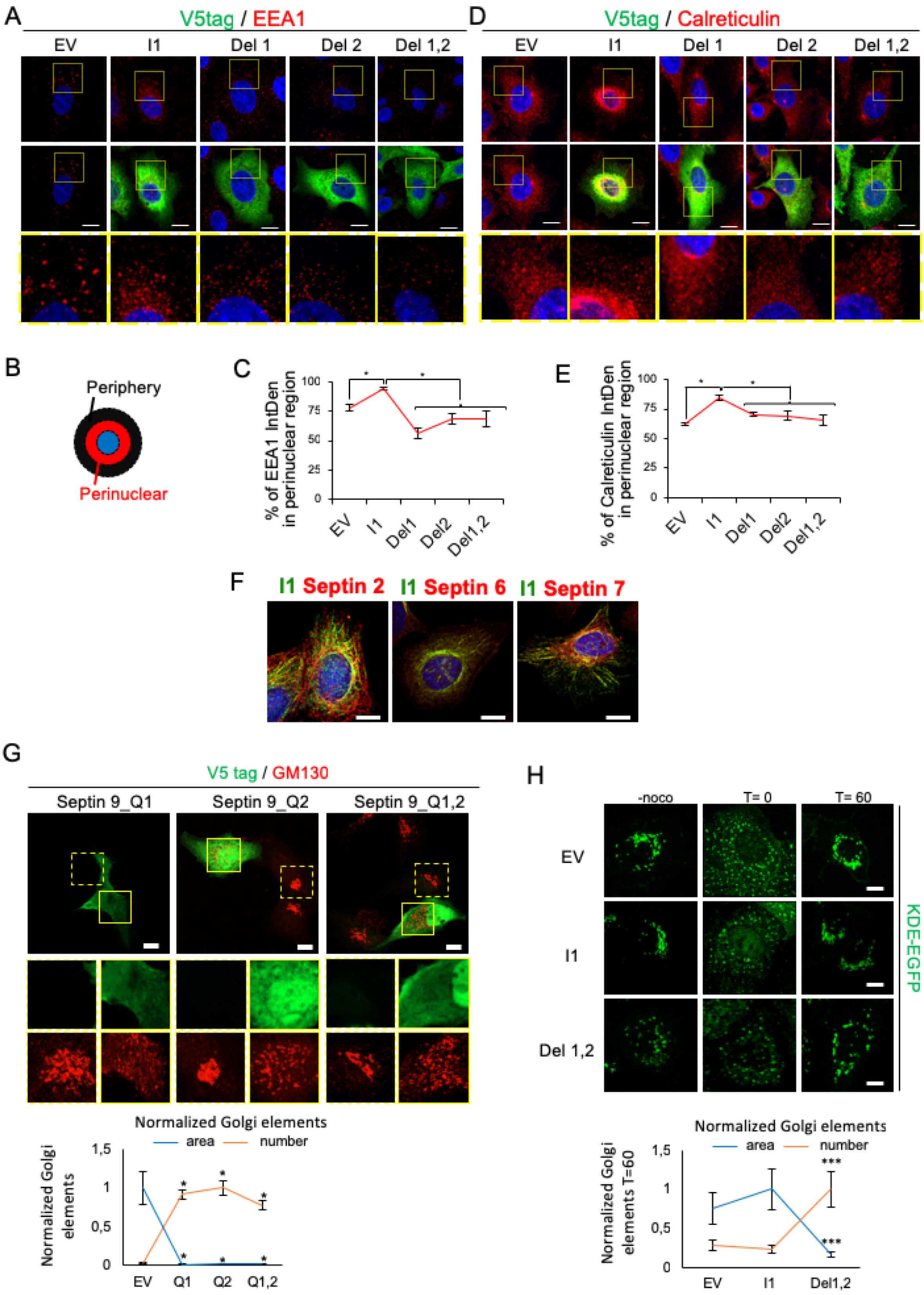


Supplementary Figure5 **(Related to Figure 5)**

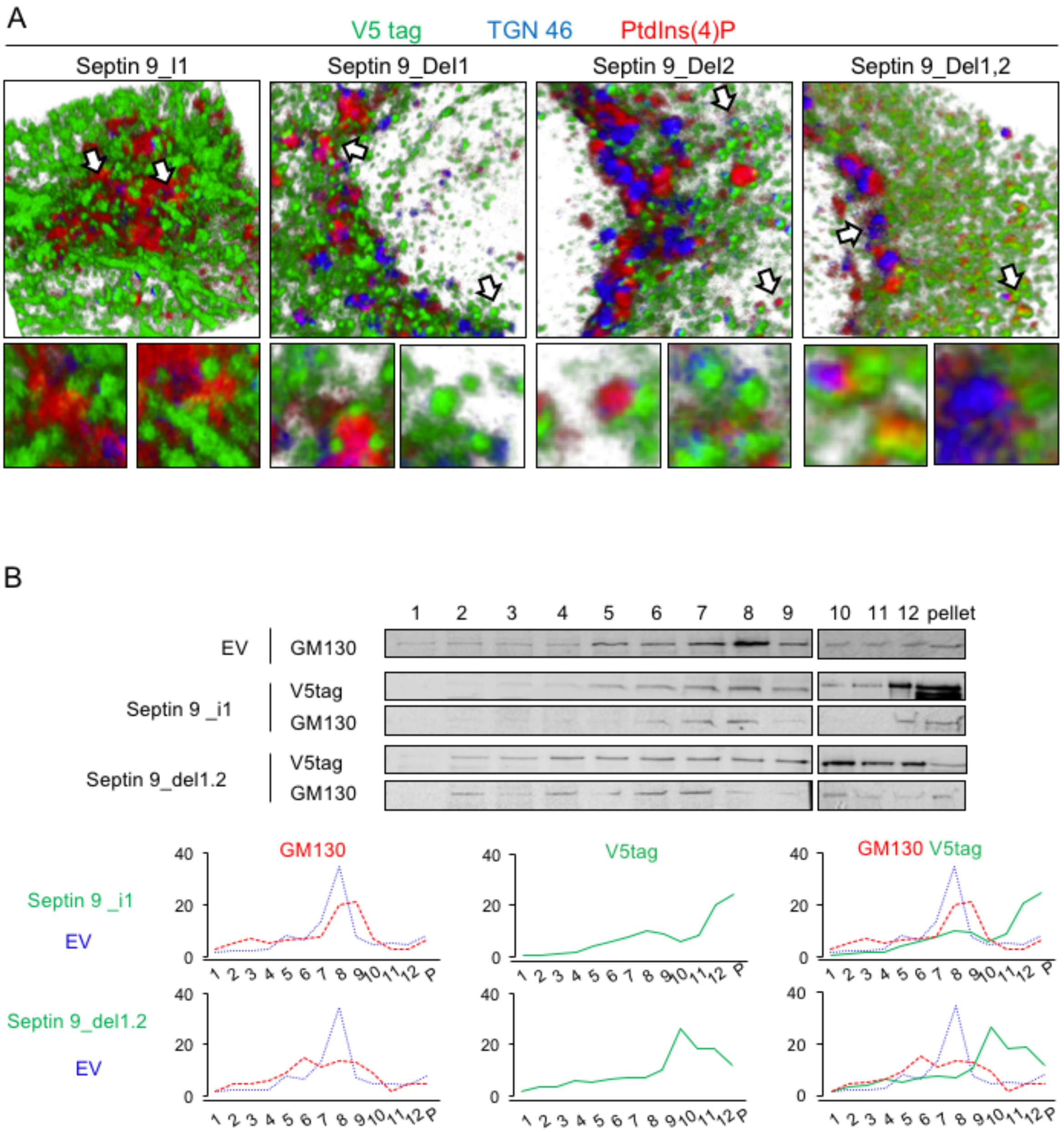


Supplementary Figure 6 **(Related to Figure 6)**

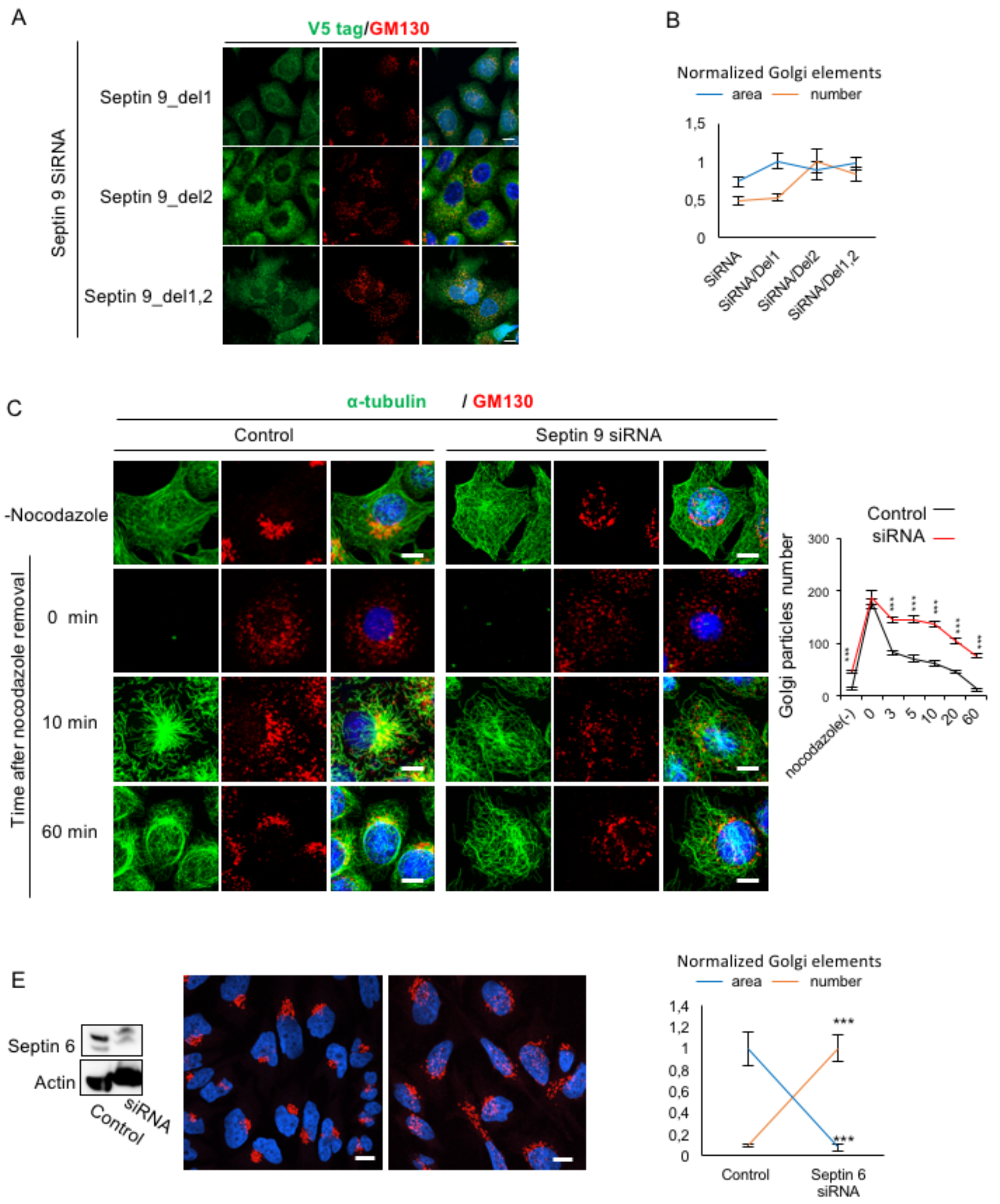


Supplementary Figure 7
**(Related to Figure 7)**

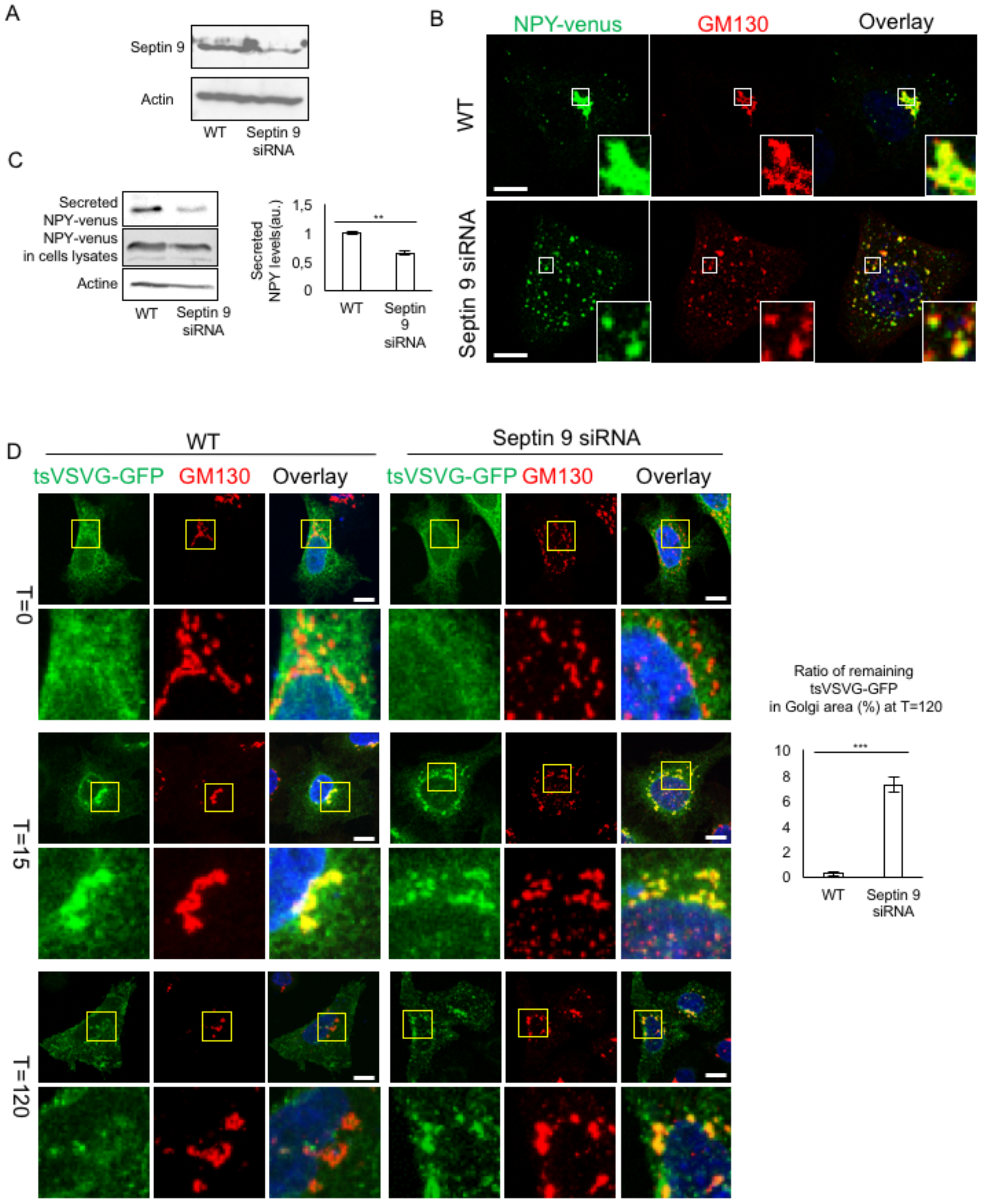


Supplementary Figure 8**(Related to Figure 8)**

#### Supplementary Figure legends

**Figure S1: Both PBs are required for septin 9_i1 assembly and the 21 residues towards N-terminal of PB1 folds in α-helix (Related to Figure 1)**

A. Multiple alignment of septin 9_I1, septin 9_del1, septin 9_del2 and septin 9_del1.2 at polybasic domain 1(PB1), G-binding domain motifs G1, G3, G4, polybasic domain 2 (PB2).
B. Coomassie blue stained SDS–PAGE gel of purified septin 9_i1 and septin 9_del1, septin 9_del2 and septin 9_del1,2.
C. Western blot of native septin 9-I1 purified protein. In the M lane Conalbumine 75kDa and Ovalbumine 45kDa were separated as reference markers of the molecular weight.
D. Analysis of secondary structure of the N-terminal region peptide computed during 500 ns of MD simulations performed in triplicate (simulation 1 to 3).

**Figure S2: PBs are required for septin 9_i1 PIs specific interaction and membranes form recognition (Related to Figure 2)**

A. Three 100 ns long unbiased molecular dynamics simulating the interaction between human septin 9 monomer and a DOPC membrane. For each simulation (MD1, MD2 and MD3) the average of contacts with the membrane atoms that appeared during these simulations were counted and are shown as bar graphs. The right-side images show the conformation of the system at the beginning of the simulation and after the end of the simulation.
B. Homology modeling structural model of septin 9 monomer showing PB1 and PB2.
C. Homology modeling structural model of the septin 9 complex G9NC/NC9G. The two molecules of septin 9 on either side of the NC interface are shown in green, and their encompassed PB1 and PB2 are shown in magenta and blue, respectively. Squares indicate PB1 and PB2 shown at a higher magnification below. The residues for PB1 and PB2 are labeled. Green arrows indicate the salt bridges between PBs and neighboring septin 9.

D. Bar graph shows protein sensibility to curvature (ratio of pound prtein with small liposomes to that of big liposomes) dashed line indicate the 1value.
E. Left, western blots of the top and bottom fractions of septin 9_del1 (Del1) and septin 9_del2 (Del2) subjected to a liposome flotation assay; the arrow indicates the band corresponding to Septin9_i1 V5 tagged (68 kDa) further analyzed. Right, bar graph shows the percentage of protein in the top fraction (bound protein) from the analysis of the blots resulting from the liposome flotation assay.

**Figure S3: Human septins have putative PB-associated amphipathic helices (Related to Figure 3)**

A. Multiple alignments of 25 residues precedent to PB1 and PB2 in human septins. The sequences of the predicted amphipathic helices (AH) associated to PBs are highlighted in yellow.
B. Bar graph representing the hydrophobic moment generated by HeliQuest of the amphipathic helices highlighted in A.
C. Bar graph representing the hydrophobicity value generated by HeliQuest of the amphipathic helices highlighted in A.
D. Bar graph representing the net charge z value generated by HeliQuest of the amphipathic helices highlighted in A.

**Figure S4: Substitutional mutations of PBs domains have a similar effect to that of deletion mutation on septins filaments. (Related to Figure4)**

A. Huh7,5 cells transfected with septin 9_Q1 (Q1), septin 9_Q2 (Q2) or septin 9_Q1,2 (Q1,2) for 48h, then fixed and stained for V5 tag (green), endogenous septin 2 (red) and endogenous septin 9 (grey). (*) indicates a low expression or not transfected cell and (0) indicates a transfected cell. Squares indicate the area shown at higher magnification to right.

B. Bar graph presenting the number of the filament structures of endogenous septin 9 and septin 2.

C. Bar graph representing the percentage of cells containing filament structures of endogenous septin 9 and septin 2.

D. Huh7,5 cells transfected with septin 9_i1 (I1), septin 9_R289A (R289A) or septin 9_R289/290A (R289/290A) for 48h, then fixed and stained for V5 tag (red), endogenous septin 2 (green) (left) and endogenous septin 9 (green) (right). (*) indicates a low expression or not transfected cell and (0) indicates a transfected cell. Squares indicate the area shown at higher magnification to the right.

E. Western blot analysis of endogenous septin 9 in huh7,5 cells transfected with septin 9_i1, septin 9_del1, septin 9_del2 septin 9_del1,2, septin 9_Q1 (Q1), septin 9_Q2 (Q2), septin 9_Q1,2 (Q1,2), septin 9_R289A (R289A) or septin 9_R289/290A (R289/290A). The tow detected bands (75kDa, 55kDa) by the antibody were presented.

**Figure S5: Mutated septin 9 i1 are incapable of having the effect of septin 9_i1 on Golgi, ER and EE compartments. (Related to Figure 5)**

A. HeLa cells transfected with either an empty vector (EV), septin 9_i1 (I1), septin 9_del1 (Del1), septin 9_del2 (Del2) or septin 9_del1,2 (Del1,2) for 48h, before being fixed and stained for EEA1 (red) and V5tag (green). The dotted square indicates the area shown at higher magnification below. Scale bar: 10μm.

B. Representation of the peripheral (black) and perinuclear (red) regions of the cell.

C. Line graph representing the percentage of EEA1 in the perinuclear region from two experiments performed as described in a. The data are shown as mean ± SEM from 10 cells under each condition.
D. Cells transfected as described in (A) were stained for calreticulin (red) and V5tag (green). The dotted square indicates the area shown at higher magnification below. Scale bar: 10µm.
E. Bar graph representing the percentage of calreticulin in the perinuclear region from two experiments performed as described in a. The data is shown as mean ± SEM of 10 cells under each condition from two independent experiments.
F. HeLa cells transfected with septin 9_i1 for 48h then fixed and stained for V5tag (green) and septin 2 or septin 6 or septin 7 in red. Scale bar: 10µm.
G. Huh7,5 cells transfected with septin 9_Q1, septin 9_Q2, septin 9_Q1,2 for 48h the fixed and stained for GM130 (red) and V5tag (green). Scale bar 10. Line graph below representing normalized Golgi elements area and number. 10 cells were analyzed form tow independent experiments.
H. MDCK stably transfected with either EV, septin 9_i1 (I1) or septin_9 del1.2 (Del1,2) were transfected with KDE-GFP for 24h to visualize the Golgi. Cells were subjected to a nocodazole washout experiment. Scale bar: 10µm. Line graph below presents the normalized Golgi elements area and numbers calculated from 10 cells after 60 minutes of nocodazole removal in two independent experiments. Student's t-test was used in C, E, G and I: *P<0.05, **P<0.001, ***P<0.0001.

**Figure S6: Septin 9 is dispensable for PtdIns4P enrichment on Golgi and their PBs are required for specific recruitment to Golgi ((Related to Figure 6)**

A. Images in Figure 6 panel (A) shown in 3D reconstruction with arrows indicating the area shown below at higher magnification. The zoomed in regions show example cases: in the septin mutants (green), the protein's signal either does not colocalizes with any of the PtdIns4P or GM130 signals (in red and blue respectively), or colocalize with only one of them, or with both.

B. MDCK EV, septin 9_i1 (I1) and septin 9_del1.2 (Del1,2) stably transfected cells were grown for 48h before being subjected to a subcellular fractionation assay and analyzed with Western blot for V5tag and GM130. The line graphs below show the densitometry analysis of the presented Western blots.

**Figure S7: Septin 9 is required for Golgi assembly (Related to Figure 7)**

A. septin 9 siRNA cells were transfected with septin 9_del1 (Del1), septin 9_del2 (Del2) or septin 9_del1,2 (Del1,2) for 48h and stained for GM130 (red) V5tag (green). Scale bar: 10µm.
B. Line graph representing the normalized area and number of Golgi elements in 15 cells from two independent experiments.
C. Septin 9siRNA and control cells were treated with nocodazole for 1 hour at 37°C and placed on ice for 2 hours. The cells were washed five times with ice-cold culture medium to remove the nocodazole and then moved to medium at 37°C for the time indicated in the Figure. The cells were then extracted, fixed and stained for αtubulin (green) and GM130 (red). Scale bar: 10µm. Line graph presenting the number of Golgi elements. Values are mean ± SEM of 30 cells from two independent experiments. Student's t-test was used ***P<0.0001.
D. HeLa cells transfected with septin 2 siRNA or septin 6 siRNA for 48h were analyzed by western blot (left) and confocal microscopy for GM130 (red) (middle). Scale bar: 10µm. Line graph presenting Golgi element's area and size. Values are mean ± SEM from 15 cells under each condition. Student's t-test was used ***P<0.0001.

**Figure S8. Septin 9 depletion affects Golgi morphology and intracellular transport (Related to Figure 7)**

A. Immunoblot of septin 9 in septin 9 siRNA and control cells.

B. Control and septin 9 siRNA cells were transfected with NPY-venus for 24h prior to fixing and staining for GM130 (Red). Scale bar: 10µm.

C. Control and septin 9 siRNA cells were transfected with NPY-venus and incubated with the same culture volume for 24h. NPY-venus was analyzed by Western blotting in the culture medium and cellular lyses. The bar graph to the right representing the mean ± sem from three independent experiments. *P<0.05, (Student's t-test).

D. Septin 9 siRNA and control cells were transfected with tsVSVG-GFP and incubated at 37 °C for 3h and then at 40°C for a further 16h prior to being incubated at 32°C for the indicated time in the presence of 50 µg/ml cycloheximide. After incubation, the cells were fixed and stained for GM130 (red). Scale bar: 10µm. Bar graph representing the remaining tsVSVG-GFP in the Golgi area after 120 minutes of incubation at 32°C. N= 10 cells from two independent experiments.

### Experimental Procedures

#### The molecular dynamics simulations

The monomer of septin 9 (Figure 1C) was built, using Coot program (Emsley et al., 2010). by adding the missing residues from chain A of the crystal structure 5cyo (which has the less number of missing residues of all solved structures for septin 9), completing the sequence from residue 275 to 563. Missing side chains were also added and we made a 1 ns NPT equilibration restraining all main chain atoms but the ones from the added residues (plus direct neighbor residues) slowly increasing the temperature from 0 to 310K. The entire structure was then equilibrated in the same manner prior to the 100 ns MD (Figure S2A).

The dimer of septin 9 (Figure 1E) was built using the above-mentioned conformation of septin 9 monomer after the missing residues were equilibrated, considering only residues from 294 to 563. Two of these conformations were superposed to the crystal structure 5cyp at the NC interface. A pre-folded N-terminal region, comprising residues 275 to 295, was added to each chain in two different orientations: one by superposing the residues 294 and 295 from the two structures and the other by superposition with the PB1 residues of the crystal structure of septin 3 (PDB code 4z54). This pre-folded N-terminal region was obtained after a 500 ns MD simulation (Figure S1D), performed in triplicate (two of them with a amide at the C-terminal), which started from a stretched random conformation of this residues manually built with Coot program (Emsley et al., 2010).

The MDs (Figure S2A) were performed with the Gromacs program (Abraham et al., 2015) using the united-atom force field gromos54a7 (Schmid et al., 2011) and a leap-frog algorithm for integrating Newton's equations of motion. Explicit solvation was used with SPC/E water model (Berendsen HJC et al.) and the DOPC membrane coordinates were obtained from the Gromacs website and the topology from Peter Tieleman's group (Tieleman et al., 2000). Periodic boundary conditions were used through all the simulations, with the long-range electrostatic interactions treated by particle mesh Ewald method (Darden et al.). The short-range non-bonded interactions were amputated with 10 Å cutoff. Bond lengths involving H atoms were constrained using the Linear Constraint Solver (LINCS) algorithm (Hess et al.). The temperature was controlled by Nosé-Hoover thermostat (Cheng and Merz, 1996), while the pressure coupling was maintained isotropic in the axes parallel to the membrane plane and different in the normal direction with the Parrinello-Rahman barostat (Parrinello and Rahman, 1981). For each simulation (MD1, MD2 and MD3) (Figure S2A) the average of contacts with the membrane atoms that appeared during these simulations were counted and are shown as bar graphs: MD1 and MD3 have the same starting protein conformation but MD1 has a lower protein velocity; MD2 has a different starting protein conformation than MD1 and MD3 but has the same initial velocity as MD3. We considered a contact whenever a protein atom was less than 4Å from a lipid atom –non-hydrogen atoms only. In MD1, the average was calculated for the entire period of 100 ns, and the conformations were sampled each 10 ps.

**Homology modeling**

**Rebuilding the structure of septin 3 (59-350)**

The crystal model of septin 3 comprises residues 59-350 of human neuronal-specific septin 3 in a complex with GDP (Macedo et al., 2013). The ordered part of this construct harbors 75% sequence identity with residues 297-565 of human septin 9 (isoform_i1; 586 residues). It starts immediately after PB1 ($^{289}$RRKAMK$^{294}$) and contains PB2 ($^{399}$RKKR$^{402}$).

We carefully completed and rebuilt the septin 3 GTPase domain G dimer available (PDB code 3SOP) using Coot (Emsley et al., 2010) and phenix.refine (Adams et al., 2010). As well as correcting a few local errors, we were able to reliably model all the missing loops, except for the N-terminus of switch I (corresponding to septin 9 residues 323-333), and most of the missing sidechains. Sidechains for which no density was apparent were still included in the most likely rotamer, given the chemical environment. The final structure comprised the counterparts of septin 9 residues 297-565 (less residues 323-333) and was refined to R_work = 0.2039, R_free = 0.2612, from the initial values of R_work = 0.2585, R_free = 0.2780.

**Homology modeling using the Rosetta of septin 9 297-565 4-molecule filament**

A septin 3, 4-molecule filament with a single NC interface was generated by means of crystallographic symmetry. The filament model of septin 9 297-565 was built using Rosetta 3.5 (Das and Baker, 2008) according to the following steps: 1) the RosettaCM comparative modelling pipeline (Song et al., 2013) was used to build a monomeric septin 9 model (Figure S2B), taking a septin 3 monomer (residues 59-350) as the template; 2) the Rosetta relax protocol was used first of all to refine the NC interface under C2 symmetry constraints, while regions distant from the NC interface remained fixed; 3) the G interface was refined in a similar way under C2 symmetry constraints, without moving the NC interface. In particular, a dozen distance constraints were carefully introduced into this step to preserve the nucleotide binding site as in the GDP-bound state; 4) finally, GDP and Mg were copied into the monomeric model and the filament was built by symmetry (Figure S2C).

**Modeling of PB1 in the context of septin 9 4-molecule filament and membrane orientation**

The threading server I-Tasser (Yang et al., 2015) was used to add septin 9 residues 288-296 (including PB1) to the 297-565 Rosetta model. Although two orientations of PB1 are possible in the context of a septin 9 monomer, only a conformation pointing away from the NC interface is consistent with filament formation across this interface (Figure S2B). We introduced this conformation into the septin 9 tetrameric model (Figure S2C). The orientation of this filament relative to a membrane was then computed using the energy transfer method implemented in the PPM server (Lomize et al., 2012).

**PC-PE-PIP /septin 9 membrane generation and optimization**

The position predicted by the PPM server was used as input to the CHARMM-GUI (Wu et al., 2014) in order to add a 200x200 Å$^2$ lipid bilayer to the septin 9 tetramer. The lipid composition was chosen to be identical for both leaflets: 55% POPC, 25% POPE, 10% POPI5P, 5% POPI3P and 5% POPI4P. This protein-membrane system was energy-minimized, equilibrated and subjected to a 10-ns molecular dynamic simulation using GROMACS (Berendsen et al., 1995) and the CHARMM36 force field update for lipids. We used the GROMACS protocols encoded in the scripts generated by CHARMM-GUI to perform this simulation (Lee et al., 2016) (Figure 2A).

**Cell lines and culture conditions**

septin 9 HeLa SilenciX (septin 9 siRNA and control cell lines) was purchased from TiboLab. HeLa and huh7 cells were maintained in a DMEM complete media composed of Dulbecco's modified Eagle's Medium (DMEM; Invitrogen) containing 4.5 g/l glucose and supplemented with 10% heat-inactivated fetal bovine serum, 1% non-essential amino acids (GibcoBRL) and 1% penicillin/streptomycin (GibcoBRL). septin 9 siRNA and control cell lines were maintained in a DMEM complete media supplemented with hygromycin B (Invitrogen)100 µg/ml. MDCK cells were maintained in MEM complete medium composed of Minimum Essential Media (MEM; Invitrogen) supplemented with 5% heat-inactivated fetal bovine serum and 1% penicillin/streptomycin (GibcoBRL). MDCK stably transfected cells of septin 9_i1 and mutant protein were maintained in MEM complete medium supplemented with G418 (Invitrogen) at 400 µg/ml.

**Plasmids**

The cDNA of the septin 9 isoform 1 transcript within the pcDNA3.1/V5-His-TOPO vector (septin 9_i1) or within the pET21d vector (pET21d septin 9_i1), PB1-deleted septin 9 within the pcDNA3.1/V5-His-TOPO vector (septin9_del1), and the pET21d vector (pET21d septin 9_del1), as previously described (Akil et al., 2016), were used during this study. Septin 9_del2 (PB2-deleted septin 9) and septin 9_del1.2 (PB1and PB2 deleted septin 9), pcDNA septin 9_Q1 (R and K residues were substituted with Q in PB1) , pcDNA septin 9_Q2 (R and K residues were substituted with Q in PB2)  pcDNA septin 9_Q1.2 (R and K residues were substituted with Q in PB1and PB2), pcDNA septin 9_R289A (R289 of PB1 was substituted with A), pcDNA septin 9_R289/290A (R289 and R290 of PB1 was substituted with A) were generated using the QuikChange II XL Site-Directed Mutagenesis Kit (Cat#200521) from Agilent, in accordance with the manufacturer's recommendations.

Septin 9_del2 V5/His tag and septin 9_del1.2 V5/His sequences were thus amplified by PCR from their corresponding pcDNA3.1/V5-His-TOPO vectors. The resulting DNA fragments were inserted in a pET21d vector using the In-Fusion HD Cloning kit Cat#639648 pET21d (Clontech), according to the manufacturer's recommendations, in order to obtain pET21d septin 9_del2 V5/His and PET21d septin 9_del1.2 V5/His. All the primer sequences are presented in Supplementary Table 1.

KDE-GFP (Dipeptidyl peptidase IV in which the extracellular domain had been replaced by the GFP sequence to restrict protein localization to the Golgi apparatus, which we used to visualize this apparatus during live cell imaging experiments) was a gift from Professor Christian Poüs (Paris-Sud University, France). NPY-Venus and tsVSVG-GFP were a gift from Professor Bruno Goud, Institut Curie, PSL Research University. Septin 6 (h) siRNA Cat#sc40938, septin 2 (h) siRNA Cat#40937 were from santacruz biotechnology.

**Chemicals:**

PIP strip Cat#P-6001, PtdIns4P diC8 Cat#P4008 were purchased from Echelon; Nocodazole Cat#487928 from Calbiochem, and Albumin BSA FFA Cat#126575 came from Calbiochem (1,2-oleoyl-sn-glycero-3-phosphocholine (DOPC) (Cat#850375 Avanti).

**Antibodies**

Anti-septin 9 Cat#ab38314 (WB:1/500, IF:1/25), anti-septin 2 Cat#ab88657(WB:1/500, IF:1/50), anti-septin 6 Cat#ab138036 (WB:1/500, IF:1/50), Anti α-tubulin Cat#ab15246 (IF:1/100), anti-giantin Cat#ab37266 (IF:1/100), mouse and rabbit anti-V5 tag Cat#ab27671 (WB:1/1,000, IF:1/400), Cat#ab9116 (WB:1/1,000, IF:1/400), FITC conjugated Cat#ab1274 (IF: 1/400) , anti-EEA1 Cat#ab2900 (IF:1/100) and anti-calreticulin Cat#ab2907 (IF:1/100) were obtained from Abcam; anti βtubulin Cat# T4026 (IF:1/100) came from Sigma-Aldrich;

anti-Actin Cat#sc-1616 (WB:1/1,000), anti-GM130 Cat# 610822 were sourced from BD Transduction Laboratories, anti-PtdIns(4)P Cat#Z-P004 (IF:1/100) was obtained from Echelon Biosciences, and anti-TGN46 Cat#NB110-62093 from Novus.

Secondary antibodies: Anti-mouse IgG-HRP and anti-rabbit IgG-HRP came from GE Healthcare (WB:1/1,000). Anti-goat IgG-HRP Cat#sc-2020 (WB: 1/1,000) came from Santa Cruz. Alexa Fluor 633 Cat#A21136, A21070 and A21082 (IF: 1/100), Alexa Fluor 568 Cat#A11004, A11011, A11057 and A21099 (IF:1/100), Alexa Fluor 488 Cat#A11001, A21206 and A21202 (IF:1/100), and nuclei stained with Hoechst Cat#H21486 (IF:1/5,000) were purchased from Life Technology.

**Immunofluorescence staining:**

Cells were grown on coverslips, fixed with 4% paraformaldehyde for 20 min. and permeabilized for 20 min. at 37°C using a permeabilizing buffer (PFS): DPBS containing saponin (Cat#10294440 Fisher Scientific) 0.025% $m.v^{-1}$ and gelatin from cold water fish skin (Cat#G7041 Sigma 0.7% $m.v^{-1}$). The cells were then incubated with primary antibody for 2 h and washed three times for 5 min. with PFS before being incubated with the appropriate secondary antibodies or with the dye for 90 min. The coverslips were mounted using Prolong Gold (Cat#P36934, Invitrogen).

**Image acquisition and analysis**

Images acquired with a Leica TCS SP5 AOBS tandem confocal microscope were analyzed using the Icy bioimage analysis software for 3D reconstruction. For co-localization analysis, images were treated with ImageJ software, and the ‘Intensity Correlation Analysis’ plug-in was used to generate Pearson's correlation coefficient (Rr) values which ranged from −1 (perfect exclusion) to +1 (perfect correlation).

To determine the distributions of EEA1 and calreticulin in cells, the ‘Radial profile’ plug-in was used. For analysis, a circle was defined at the periphery of each cell and the plug-in produced a profile plot of normalized integrated intensities around concentric circles as a function of distance from a point in the image, considered here as the center of the cell. The concentric circles were assembled in three circle bands, the first corresponding to the area of the nuclei and the rest corresponding to the cytoplasm which was divided in two equal bands (the band near the nucleus being considered as ‘perinuclear’ and the other as the ‘periphery’). The intensity in each band was calculated from the total integrated intensities around the concentric circles present in the band.

To calculate the size and number of Golgi elements, images obtained by confocal microscopy were processed by background subtraction and standardized thresholding (default). The cell ROI was obtained using a freehand selection tool, and then the size and number of Golgi elements were determined by ImageJ particle analysis function (particle areas smaller than 0.01 μm2 were excluded).

The Golgi compactness index, which determines a dimensionless circularity of Golgi elements, was computed according to the formula 4 (sum (areas)/(sum(perimeters))^2). The values of this index ranged from 1 (perfect compactness) to 0 (perfect fragmentation)(Bard et al., 2003).

For endogenous septin 9 analysis in the filaments of high and low septin 9_i1 expressed cells (Figure 4 panel B and C) we have calculated the ratio of septin 9 intensity in a filament of septin 9_i1 low expression to that of high expression. To ensure that we compare a similar filament in size we have calculated the ratio of septin 9 in a filament with similar septin 2 intensity for that we have the ratio of septin 2 at a value of almost 1 and we observed how the septin 9 ratio will vary.

**Liposome floatation assay:**

Liposomes were prepared as follows. Two quantities of 1.5µM of DOPC were dissolved in chloroform and dried to a film under nitrogen gas. 1ml of HKM buffer (50 mM Hepes, 120 mM K acetate, and 1 mM $MgCl_2$, pH 7,4.) was added to the first quantity and vortexed to produce the control liposomes (PtdIns4P (-)). The same volume of HKM buffer containing PtdIns4P at 50µM was added to the second quantity and vortexed to produce phospoinositide 4 monophosphate containing liposomes (PtdIns4P (+)) at 3.33% (PtdIns4P to DOPC molar percentage). All the liposomes thus produced were then subjected to seven freeze–thaw cycles and considered as large liposomes. 500µl of PtdIns4P(-) and PtdIns4P(+) large liposomes were then sonicated for 30 seconds to generate the small liposomes.

septin 9-i1 and mutant proteins were added to 200 µl of liposome to a final concentration of 0,1µM, were mixed with an equal volume of sucrose at 75% and then layered on the bottom of a 1-ml thick wall ultracentrifugation tube. 200µl of 20% sucrose and 200 µl of 10% sucrose and 200 µl of HKM buffer were then layered successively. Each tube was subjected to centrifugation at 30,000 rpm using a SW60 rotor (Beckman) for one hour at 4°C. The liposomes were collected from the top 100 µl of the gradient (top fraction) and 100 µl were collected from the bottom of the tub (bottom fraction). The collected fractions were then analyzed using Western blot. The sum of the three bands detected by the V5 antibody in each lane was used to generate the presented results in Figure 2 and Figure S2. To analyze the sensitivity of the protein to bind the PtdIns4P we have calculated the ratio of detected protein in the top fraction of PtdIns4P(+) to that of PtdIns4P(-) in the case of big liposomes and small liposomes and we have call it protein sensitivity to PtdIns4P. To analyze the sensitivity of the protein to bind the small liposomes we have calculated the ratio of detected protein in the top fraction of the small liposomes to that bound to big liposomes in the case of PtdIns4P(+) liposomes and PtdIns4P(-) liposomes and we have call it protein sensitivity to curvature.

**Subcellular fractionation assay**

Confluent monolayers of cells were placed on ice, washed twice with ice-cold PBS at pH 7.4, and then 10 mM Tris/HCl (pH 7.4) buffer was added for 1min. The cells were scraped into a homogenization buffer comprising 10 mM Tris/HCl, 1 mM EGTA, 0.5 mM EDTA and 0.25 M sucrose, at pH 7.4, which also contained Complete$^{TM}$ protease inhibitors. The result was homogenized with ten strokes of a loose-fitting Dounce homogenizer. All subsequent steps were carried out at 4°C. Post-nuclear supernatants (PNS) were obtained by centrifugation for 10 min at 1000 g. PNS (2 ml) were layered onto a sucrose gradient [successive layers of 1 ml of sucrose at 40% (w/v), 1 ml at 30%, 2ml at 25%, 2ml at 20%, 2ml at 15% and 2 ml at 10%]. The gradients were centrifuged for at least 16 h at 175,000 rpm. After centrifugation, twelve 1-ml fractions were harvested, starting at the top of the gradient. The pellet was re-suspended in 1 ml 0.25 M sucrose (Waugh et al., 2003) (Figure S6B). In Figure 7B, thirteen fractions were collected and the pellet was designated as fraction 14.

**Protein production and purification**

*Escherichia coli* BL21(DE3) Rosetta cells were transformed using pET21d septin 9_i1 V5/His, PET21d septin 9_del1 V5/His, PET21d septin 9_del2 and PET21d septin 9_del1,2 V5/His V5/His vectors, and were all incubated with a culture medium containing 100 µg/mL ampicillin and 34 µg/mL chloramphenicol. Cells from a single colony were used to seed an overnight 20 ml pre-culture of LB medium. 1 L LB medium cultures from these pre-cultures were grown at 37°C under agitation to reach an optical density ($OD_{600\ nm}$) of 0.9. The temperature was lowered to 28°C and protein expression was then induced with 1mM Isopropyl β-D-1-thiogalactopyranoside (IPTG) from Sigma, for 4 hours. The cells were harvested by centrifugation and the pellet was stored at -80°C until use. The cell pellet was suspended in 15 mL of 50 mM sodium phosphate at pH 7.4, 300 mM sodium chloride, 10% glycerol, 20 mM imidazole, 0.1% Triton X-100 and one protease inhibitor cocktail EDTA-free tablet (Roche). Cell lysis was performed by sonication on ice and the cell lysate was clarified by centrifugation for 30 min. at 4°C and 40,000 rpm. After filtering the supernatant, the protein was isolated on a 1 mL metal affinity column (HisTrap HP, GE Healthcare) pre-equilibrated in 50 mM sodium phosphate pH 7.4, 300 mM sodium chloride, 10% glycerol, 20 mM imidazole and then eluted with 50 mM sodium phosphate pH 7.4, 300 mM sodium chloride and 300 mM Imidazole. The fractions containing the protein of interest were further purified by cation exchange chromatography. The pooled fractions were diluted at a ratio of 1:5 to obtain a final solution of 30 mM Tris pH8, 100 mM NaCl, 1 mM EDTA and then incubated with 1 mL of a strong cation exchange resin (Macro-Prep 25S, BioRad). The resin was packed in a column and eluted using a 100 to 600 mM NaCl linear gradient. Fractions containing a majority of protein were obtained at around 350 mM NaCl. The proteins were then flash-frozen and stored at -80°C.

**Determination of helix properties**

Helical wheels were generated using the Heliquest server (Gautier et al., 2008) http://heliquest.ipmc.cnrs.fr/

**PIP strip overlay assay**

PIP Strip membranes (Echelon Biosciences) were blocked with 3% BSA FFA dissolved in phosphate-buffered saline (PBS) containing 0.1% Tween 20 (3% BSA FFA PBS-T) at room temperature for 60 min., then incubated overnight at 4°C with the same buffer containing the purified protein of interest at a concentration of 0.5 µg ml−1, or the purified V5 tag peptide at an equivalent molar concentration, to act as a control. The membranes were then washed, and bound proteins were detected using a suitable antibody.

**tsVSVG secretion assay:**

Sept9siRNA and control cells were transfected with tsVSVG-GFP and incubated at 37°C for 3h then at 40°C for a further 16h before being incubated at 32°C for the indicated time in the presence of

50 µg/ml cycloheximide (Cat: C4859 Sigma). After incubation, cells were fixed and stained.

**Immunoblotting**

The cells were washed with ice-cold DPBS and lysed on ice using the following buffer: 20 mM Tris, HCl, 100 mM NaCl and 1% Triton X100 at PH 7.4 containing protease inhibitors (cOmplete™ ULTRA Cat#05892970001 Roche). The proteins were separated on SDS (sodium dodecyl sulfate) polyacrylamide gel and electro-transferred onto nitrocellulose membranes. After transfer, the membranes were saturated in DPBS containing 0.1% Tween 20 and 5% milk. Primary antibodies were added overnight at 4°C or for 2 h at room temperature, depending on the antibody. The membranes were then washed with DPBS and incubated for 1 h at room temperature with appropriate secondary antibodies coupled with peroxidase. The ECL plus kit (Cat#32132), SuperSignal™ West Femto Maximum Sensitivity Substrate (Cat#34095) from Thermo Scientific were used for protein detection. Chemiluminescent signals were detected by the G:BOX Chemi Fluorescent & Chemiluminescent Imaging System from SYNGENE. The blots were quantified using ImageJ software. For native purified protein separation (Figure S1C). NativePAGE™ 3-12% Bis-Tris Protein Gels, cat# BN1001BOX from Thermo Fisher scientific were used then the protein were electro-transfered onto PVDF membrane then treated as described above. For NPY analysis in culture medium (Figure S8C), 40µl of culture medium were separated by SDS-PAGE and then processed as indicated above.

**Nocodazole washout assay**

Cells were grown on 12-mm glass coverslips and allowed to attach overnight before being incubated with a culture medium containing nocodazole (2.5µg/ml) for 1 hour at 37°C, then for 2 hours on ice. The cells were rinsed five times on ice using ice-cold medium in order to remove the nocodazole and were then moved to a pre-warmed medium (at 37°C) (time 0) and incubated at that temperature for the indicated time before being fixed and stained.

To detect microtubules in the washout experiments, soluble tubulin was eliminated first of all in order to reduce background fluorescence. For this purpose, cells were extracted with MT-stabilizing buffer (80 mM Pipes, 1 mM MgCl2, 2 mM EGTA; pH 6.9) containing 0.1% Triton X-100 (37°C, 30 seconds), and then washed twice with Triton-free buffer at room temperature before being fixed with methanol at -20°C(Poüs et al., 1998).

Imaging the assembly of Golgi apparatus after the removal of nocodazole was achieved using an AXIO-OBSERVER Z1 – COLIBRI® – TIRF 3 VIDEOMICROSCOPE with 6 s between frames, and an exposure of 200 ms (inverted images) (Fig. 3b).

**Statistical analyses**

Unpaired Student's t-tests were performed and statistical significance was determined at *P <0.05, ** P <0.001 and ***P < 0.0001.

### Table S1

| | |
|---|---|
| Pet21d septin 9_i1-F | CTCCGTCGACAAGCTATGAAGAAGTCTTACTCAGGAGG |
| Pet21d septin 9_i1-R | GGTGGTGGTGCTCGATCAATGGTGATGGTGATGAT |
| septin 9_del1 F | CATCCTGGAGCAGATGCAGGGCTTCGAGTTCA |
| septin 9_del1 R | TGAACTCGAAGCCCTGCATCTGCTCCAGGATG |
| septin 9_del2 F | GTCAACATCAACATCCCGGACACCCG |

| | |
|---|---|
| septin 9_del2R | CGGGTGTCCGGGATGTTGATGTTGAC |
| septin 9_Q1F | TCCTGGAGCAGATGCAGCAGCAGGCCATGCAGCAGGGCTTCGAGT |
| septin 9_Q1R | ACTCGAAGCCCTGCTGCATGGCCTGCTGCTGCATCTGCTCCAGGA |
| septin 9_Q2F | GAGGAGGTCAACATCAACCAGCAGCAGCAGATCCCGGACACCCGCGTCC |
| septin 9_Q2R | GGACGCGGGTGTCCGGGATCTGCTGCTGCTGGTTGATGTTGACCTCCTC |
| septin 9_R289AF | CCTGGAGCAGATGGCCCGGAAGGCCATG |
| septin 9_R289AR | CATGGCCTTCCGGGCCATCTGCTCCAGG |
| septin 9_R289/290AF | TGGAGCAGATGGCCGCGAAGGCCATGAAGC |
| septin 9_R289/290AR | GCTTCATGGCCTTCGCGGCCATCTGCTCCA |
